# Cortical GABAergic inhibition dynamics around hippocampal sharp-wave ripples

**DOI:** 10.1101/2025.08.17.670701

**Authors:** Edris Rezaei, Setare Tohidi, Mojtaba Nazari, Bruce L. McNaughton, Majid H Mohajerani

**Author notes:** Corresponding author to: Edris Rezaei.

## Abstract

Hippocampal sharp-wave ripples (SWRs) coordinate hippocampal–neocortical interactions for memory consolidation, yet how cortical GABA signaling is organized around SWRs remains unclear. Here we approach this problem by combining wide-field mesoscale imaging of extracellular GABA using iGABASnFR2 with simultaneous dorsal CA1 electrophysiology and sleep-state monitoring in mice, enabling GABA dynamics to be mapped across 17 cortical regions during natural sleep and wakefulness. Using ripple-triggered activity mapping and singular value decomposition, we identified a cortex-wide GABA response consisting of a dominant global component and regionally structured components that were reconfigured across brain states. Across cortical subnetworks, SWRs were associated with a reduction in GABA signaling followed by widespread activation, with both components enhanced during NREM sleep. During NREM sleep, GABA responses emerged earliest and most strongly in the retrosplenial and other medial cortical regions before progressing laterally. During wakefulness, responses were faster, preferentially recruited lateral sensory regions and progressed towards medial cortex. Transitions between NREM sleep, REM sleep and wakefulness were also accompanied by distinct changes in GABA signaling and interregional network organization. Our findings suggest a model in which hippocampal SWRs recruit a shared cortex-wide GABA response whose regional expression and direction of propagation are reconfigured by brain state, providing a dynamic inhibitory framework for regulating hippocampal–neocortical communication.

## Introduction

Cortical function is result from the coordinated interaction between excitation and inhibition that together determines the dynamics of neural circuits and behavior ^1^. Excitatory neurons generate most cortical output and communicate long distance between brain regions^2^. This excitatory flow must be modulated to ensure stable network, accurate information processing, and adaptive behavioral output ^3^. GABAergic interneurons, about 20–30% of the cortical neurons, regulating circuit excitability, shaping neuronal response, and organizing rhythmic network activity^4–5^. Cortical GABAergic interneurons are broadly classified into three major subtypes based on molecular expression: parvalbumin-positive (PV+), somatostatin-positive (SST+), and vasoactive intestinal peptide–positive (VIP+) interneurons^5–7^. PV+ interneurons target the peri-somatic region of pyramidal neurons and exhibit fast-spiking firing properties, enabling them to provide powerful feedforward and feedback inhibition and to synchronize network activity. PV+ interneurons suppress pyramidal cell firing and generating gamma-frequency oscillations ^9–10^. SST+ interneurons, in contrast, inhibit distal dendrites of pyramidal neurons and provide feedback inhibition that regulates dendritic integration of top-down and long-range inputs ^11–13^. VIP+ interneurons inhibit other interneurons, particularly SST+ cells, thereby recruiting disinhibitory circuits that sustain pyramidal neuron output during behavioral states such as attention and changes in arousal^14–16^. GABAergic interneurons shape synaptic plasticity, tune excitatory drive, and synchronize communication across distributed cortical networks. The coordination between excitation and inhibition dynamically varies with behavioral state, including sleep, wakefulness, attention, and locomotion. Disruption of GABAergic interneurons has been implicated in a broad spectrum of neuropsychiatric disorders, including epilepsy, autism spectrum disorder, and schizophrenia^1,6,17,18^.

Earlier electrophysiology established that hippocampal sharp-wave ripples are temporally coordinated with neocortical network activity during sleep. Simultaneous recordings showed that neocortical slow oscillations and spindles often precede and modulate ripple generation in the hippocampus, suggesting that cortical activity can help organize hippocampal events^19^. Other studies showed that hippocampal sharp-wave ripple bursts occurred around the time when the cortex entered an active “up-state” and could influence cortical population activity, suggesting that hippocampal activity can also shape cortical dynamics^20^. Together, these studies showed that the hippocampus and neocortex interact in both directions during sleep. This coordinated hippocampal–cortical activity is thought to support memory consolidation and provides an important electrophysiological foundation for later optical imaging studies. In this context, it is important to understand how cortical GABAergic inhibition is regulated by internally generated hippocampal activity patterns, such as sharp-wave ripples. SWRs are well positioned to coordinate brain-wide communication during sleep and quiet wakefulness.

Recent studies have started to answer this question by describing excitatory cortical activity in response to hippocampal SWRs. In a study using iGluSnFR transgenic mice and voltage-sensitive dye imaging, Karimi Abadchi et al.^21–22^ demonstrated that, during NREM sleep, the SWRs precede a period of suppression in the whole cortex that is followed by sequential activation originating in medial and in visual cortices, areas involved in the default mode system. During quiet wakefulness, however, SWRs are followed by robust suppression in agranular retrosplenial cortex, with late rebound in cortical glutamatergic activity. These studies demonstrate that SWRs have organized and state specific excitatory patterns of excitation as their outcome. Cortical computation, however, is almost never regulated by excitation in isolation. Since excitatory drive is generally followed or accompanied by inhibition ^1,6^ the state-dependent patterns of excitation during SWRs thus bring to the forefront the question of the corresponding inhibitory patterns. Thus, although cortical excitatory responses to SWRs have been well described, the inhibitory component of these cortical responses remains largely unexplored. Here, we aimed to address this gap by combining wide-field imaging of extracellular GABA using iGABASnFR2^23^ with hippocampal electrophysiology. Because the sensor was expressed mainly in cortical excitatory neurons, this approach allowed us to track extracellular GABA dynamics across the dorsal cortex during different brain states, including NREM sleep, REM sleep, and wakefulness. These signals reflect changes in the local GABAergic inhibitory environment around cortical excitatory neurons, rather than direct postsynaptic inhibitory currents. Extracellular GABA mainly comes from synaptic release24, but it can also arise from non-synaptic sources, including astrocytes^25^. In addition, the effect of GABA can vary depending on brain state because chloride gradients change across states^26^.

We first established a baseline by characterizing spontaneous cortical GABA transients across different brain states. This allowed us to compare SWR-related GABA dynamics against the ongoing state-dependent inhibitory background. We then asked whether hippocampal SWRs shape cortical GABAergic activity in organized temporal and spatial patterns. By focusing on the inhibitory component of the cortical response to SWRs, our study aims to reveal how hippocampal activity is linked to distributed cortical inhibitory dynamics and how this organization may influence information flow across the brain.

## Materials and methods

### Animals and experimental conditions

We employed eight adult (>2 months) EMX-Cre female and male mice of the C57BL/6J strain for imaging studies. For studying natural sleep, EMX-Cre mice were injected intracranially with AAV vectors in a Cre-dependent manner to drive the expression of the genetically encoded GABA sensor iGABASnFR2 in neocortical and hippocampal excitatory neurons. This enabled optical reporting of extracellular GABA in excitatory cortical circuits. Strong expression of iGABASnFR2 in targeted brain areas was validated through brain sectioning analysis as outlined in our previous publication ^27^. Mice were group-housed (two to five mice) under a 12-hour light dark cycle and had ad libitum access to water and a standard laboratory mouse diet. For sleep recording studies, mice were singly housed after head-plate and electrode implantation surgery. All animal studies were approved by the University of Lethbridge Animal Care Committee and adhered to the Canadian Council for Animal Care guidelines.

### Viral constructs and retro-orbital injection procedure

To measure extracellular GABA dynamics mainly around cortical excitatory neurons, we used Emx1-Cre transgenic mice. In these mice, Cre recombinase is expressed mostly in excitatory pyramidal neurons of the neocortex and hippocampus^28^. Although Emx1 expression is mainly found in excitatory neurons, some expression may also occur in glial cells, including astrocytes and oligodendrocytes. These glial cells can express GABA receptors and, in some cases, release GABA^29–31^. To further restrict sensor expression to neurons, we used AAV vectors with the Synapsin promoter, which is largely inactive in astrocytes and other glial cells. Therefore, iGABASnFR2 expression was mainly neuronal. The sensor signal reflects extracellular GABA in the local environment around excitatory neurons, rather than direct postsynaptic inhibitory currents. We used two viral vectors: AAV2/PHP.N-Syn-FLEX-iGABASnFR2, which expresses the Cre-dependent GABA sensor, and AAV2/PHP.N-Syn-FLEX-cpSFGFP, which expresses a non-sensing GFP control. Both viruses were obtained from the Canadian Neurophotonics Platform Viral Vector Core (RRID: SCR_016477) and were purified to a final concentration of 1.5 × 10¹³ genome copies per milliliter (GC mL□¹). Viruses were delivered by retro-orbital intravenous injection^32^ in 6-week-old male and female mice. Mice were anesthetized with isoflurane (maintained at 1.5–2%), and Metacam (5 mg kg□¹) was given for pain relief. Before the injection, one drop of proparacaine ophthalmic anesthetic was applied to the eye. A 30-gauge needle was inserted into the medial canthus at about a 30° angle, and 1.4 × 10¹¹ genome copies of virus were injected into the retro-orbital sinus. After injection, the AAV2/PHP.N capsid crossed the blood–brain barrier and produced widespread brain expression, although expression efficiency can vary by brain region and mouse strain ^33,34^. In our previous study, we found robust and uniform iGABASnFR2 expression four weeks after injection ^27^. Therefore, all imaging and electrophysiological recordings in the present study were conducted ≥ 4 weeks after viral delivery to ensure mature expression and reliable GABA signal detection.

### Surgical procedure and electrophysiological recording setup

Cranial window and electrode implantation were performed 4 weeks after viral injection, once robust cortical expression of iGABASnFR2 had been established. The animals were given buprenorphine (0.05–0.1 mg/kg, subcutaneously) 30 minutes prior to surgery, and then isoflurane (1–2% oxygen) was administered via a nose cone. The scalp was shaved and sterilized, and the skull was covered with C&B Metabond (Parkell, Inc.), to which a custom-designed headplate was attached over the skull. A 12-mm glass coverslip (Carolina Biological Supply, Cat. No. 633005) was then cemented to establish a chronic cranial window to image at the mesoscale. The LFP recordings were obtained through teflon-coated 50 μm stainless steel wires (A-M Systems) implanted for electrophysiology. A bipolar electrode was positioned in the right hippocampus through craniotomy at approximately 2.6 mm laterally from the midline, tangent to the posterior edge of the occipital suture. The electrode was advanced at a 57° angle relative to vertical while monitoring neural signals both visually and audibly. Once optimal signal quality was achieved, the electrode was fixed in place using Krazy Glue and C&B Metabond. In addition, a bipolar EMG electrode was implanted in the nuchal muscles to record muscle tone.157 LFP signals from hippocampal electrode was amplified (Å∼1,000), bandpass filtered (0.1–10,000 Hz) using a Grass A.C. pre-amplifier (Model P511, Artisan Technology Group), and digitized at 20 kHz using a Digidata 1440 acquisition system (Molecular Devices Inc.).

### Habituation for head-restraint sleep and wakefulness experiments

They were then gradually introduced to the recording device after 7 days of recovery by positioning the animals on the recording platform. The first step consisted of letting the animals get to the platform and adapt to the environment. On subsequent days, having settled, they were positioned on head restraint in step increases in the period of head fixation, initiated at five minutes during the first day and incremented in five-minute increases to a period of one hour total.

### GABA imaging during wakefulness

Following the habituation, every animal was recorded every three days, with three to four sessions being done on each mouse. The recording was always done at the same time of the day to reduce variability and stress. Finally, the animals were put back into their home cages in the housing pod, where they rested and recovered before the subsequent recording.

### GABA imaging during natural sleep

Sleep recordings were performed after the completion of all wakefulness sessions. To optimize conditions for natural sleep under head restraint, the day before recording, the animals were moved from their home cage in the colony room to a separate room at noon. Sleep was restricted to ∼6 hours by gently stimulating the mice with a cotton-tip stick whenever they showed signs of drowsiness. The animals were then transferred to a larger cage containing new objects, such as a running wheel, Cheerios, and a water container, and left in surgical recovery where the temperature was controlled. The next day, the animals were transferred to the imaging room, and sleep recordings began at ∼7:00 AM. Once recording finished, the animals were returned to their home cages in the podding room and allowed to sleep freely for at least three days before repeating the procedure for another recording.

### GABA imaging

Imaging was undertaken with a custom macroscope comprising a front-to-front video lens setup, 8.6x8.6 mm field of view. The focal plane was set ∼0.5–1 mm below the cortical surface. Fluorescence signals were recorded with a 12-bit CCD camera (1M60 Pantera, Dalsa, Waterloo, ON) attached to an EPIX E8 frame grabber and operated by XCAP 3.8 software (EPIX, Inc., Buffalo Grove, IL), imaging at a frame rate of 80 Hz. Imaging parameters have been used in previous studies, including voltage-sensitive dye imaging ^35–37^. Sequential illumination was provided using alternating blue (473 nm) and green (530 nm) light-emitting diodes (LEDs) (Thorlabs) ^38^. Blue light (473 nm, filtered through a 467–499 nm bandpass) was employed to excite the iGABASnFR2 indicator, and green light (530 nm, filtered through a 527/42 nm bandpass) for intrinsic signal imaging of blood volume. A bandpass emission filter was placed in front of the CCD camera to facilitate selective detection of either fluorescence or reflectance signals. Blue and green LEDs were used as illumination sources. They were alternatively synchronized and frame-by-frame, utilizing transistor–transitor logic (TTL) triggering, to offer temporal control precisely to interleave acquisitions of reflectance and fluorescence images at 40 Hz per channel. Reflectance images, which may be used for corrections of blood artifacts ^39–41^, were analyzed as well within our current pipeline.

### Preprocessing

Image data were transferred into MATLAB; blue and green frames were differentiated based on reference frames to set up illumination origins. Only the region of interest (ROI) was selected utilizing a binary mask. Slow baseline drifts were eliminated utilizing a local detrending algorithm, based on a filter of the Chebyshev type. The ratio of ΔF/F□was normalized to the fluorescence signals (F) based on the following formula: ΔF/F□= (F□− F□) / F□× 100, where the detrended, curve provided an estimate of F□. To filter out high frequency noise, a low-pass finite impulse response (FIR) filter, utilizing a 6 Hz cut-off, was used on the time series of ΔF/F□. Hemodynamic artifacts in the blue channel were corrected by linear regression, using the green reflectance signal as a regressor. To denoise the data further, singular value decomposition (SVD) was applied to the corrected blue channel, and the first 300 components were used for reconstruction.

### Sleep scoring

Motion data were obtained from behavior camera (frame rate = 25HZ) by the FaceMap ^42^, which computes frame-to-frame pixel intensity differences in manually defined regions of interest (ROIs). Five ROIs—nose, whisker pad, ear, shoulder, and trunk—were selected to monitor both subtle facial expressions and body movement. Motion signals were time-locked to electrophysiological recordings. Wakefulness in this study was identified by clear motor activity in video recordings and high EMG amplitude. NREM sleep was determined by decreased EMG tone, low hippocampal theta-to-delta power ratio, and the occurrence of large irregular activity in the hippocampal LFP. REM sleep was characterized by EMG suppression, sustained hippocampal theta oscillations, and high theta-to-delta ratios. This multimodal sleep-scoring pipeline, combining motion-tracking, EMG amplitude, and hippocampal spectral features, is illustrated in **Video 1**. Across the seven animals used for wakefulness analysis, total recording time ranged from 35 to 142 minutes. On average, mice spent 70.5 ± 8.4% of the session in wakefulness, corresponding to 55.3 ± 28.8 minutes. Sleep-state analysis was performed in five animals. In these animals, mice spent 27.7 ± 6.7% of the recording time in NREM sleep, corresponding to 21.1 ± 8.4 minutes, and 3.8 ± 1.1% in REM sleep, corresponding to 2.8 ± 1.3 minutes. These sleep–wake proportions are consistent with what is typically observed during head-fixed mesoscale imaging experiments (Table 1).

**Tabel 1.**
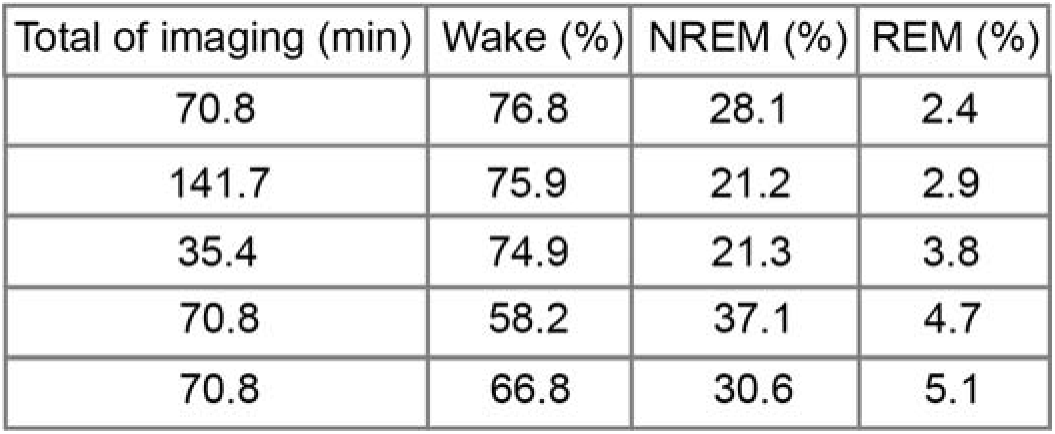
Sleep–wake distribution across animals.

### SWR detection

SWRs were detected using a modified version of the protocol described by Karimi Abadchi ^21^et al. The raw hippocampal LFP signals were first down-sampled to 2 kHz and filtered in the ripple frequency range of 110–250 Hz using a real-valued Morlet wavelet in MATLAB. The filtered ripple-band signal was then rectified and smoothed to generate a ripple-power trace. Candidate SWRs were identified when ripple power crossed a threshold defined as the baseline mean plus a multiple of the baseline standard deviation. This threshold multiplier was adjusted separately for each animal to account for differences in signal quality. For each detected event, onset and offset were defined as the times when ripple power crossed 75% of the detection threshold. Events were then filtered by duration, and events shorter than the mean duration of all detected ripples were excluded. The remaining SWRs typically lasted 50–150 ms, consistent with previously reported SWR durations in rodents ^43–44^. For each SWR, the event center was defined as the time point of the largest negative deflection, or trough, in the ripple-band signal. Events with centers occurring within 50 ms of each other were merged and treated as a single SWR.

### Normalization of peri-SWR neocortical activity using Z-scoring

To normalize peri-SWR neocortical GABA activity, random timestamps were created by shuffling the inter-SWR intervals within individual recording sessions. Imaging stacks 0.5 seconds before and 0.5 seconds after the timestamp were sampled and time-aligned for each random event. Averaging peri-event stacks across all random timestamps were then calculated to compute a baseline mean stack, as well as a parallel SD stack for standard deviation. These mean and SD stacks served as references to z-score normalization. The peri-SWR imaging stacks were then all normalized by subtraction of the baseline mean stack and division by the SD stack on a pixel-by-pixel basis to provide a z-scored representation of neocortical activity.

### SVD decomposition and component selection

For each animal, we built a peri-ripple data matrix by concatenating widefield imaging frames from −1 to +1 s around ripple centers (40 Hz; ripple center at 0 s). Non-cortical pixels were masked, and the resulting matrix was mean centered across time before performing singular value decomposition (SVD; X ≈ UΣV□). The variance explained by each component was calculated as (σ□²/∑σ□²), and the cumulative variance curve was used to determine the number of components to retain. Components up to the first point reaching ≥95% cumulative variance (median across animals: [insert value]) were kept for quantitative analyses. For visualization, we show the first five components because these components were consistent across animals and captured the main state-dependent spatial and temporal patterns. Higher-order components with lower variance were grouped together as components 2–100 and compared with the dominant global component, component 1. To show the temporal dynamics, temporal modes were averaged across ripples and plotted from −1 to +1 s relative to the ripple center. These traces were lightly smoothed using a 5-point moving average.

## Statistical Analysis

All statistical analyses were conducted using built-in functions in MATLAB.

## Data availability

All data and custom software/algorithms necessary to interpret and replicate the findings and methods of this article are available upon request.

## Results

### Experimental protocol for investigating neocortical GABA dynamics

We retro-orbitally injected AAV2/PHP.N-Syn-Flex-iGABASnFR2 or control construct (cpSFGFP) into animals (see **Figure 1A**). The configuration of our recording involved mesoscale GABA imaging, in-vivo electrophysiological recordings, and monitoring of behavior (see Figure 1B). Protocols of imaging were conducted at 80 Hz cycling blue (470 nm) and green (530 nm) LEDs alternating with a CCD camera. LFPs were recorded from the right dorsal CA1 to study hippocampal SWRs, with neck EMG and a camera for monitoring behavior. A schematic of the headplate placement and the 17 cortical regions imaged in the right hemisphere is shown in Figure 1Ci–ii. Example cortical images are shown in **Figure 1Ciii–iv**, with the dorsal retrosplenial cortex (dRSC) marked as a representative ROI. To correct for hemodynamic artifacts, GABA fluorescence signals were normalized to simultaneously recorded reflectance traces (**Figure 1Cv–vi**). Frames of images were registered to the center of SWR activity (see **Figure 1Di**), averaged and concatenated into single peri-ripple montages (see **Figure 1Di–ii**). To reveal cortical GABAergic inhibition dynamics, we provide two companion videos. **Video 2** provides spontaneous neocortical GABA dynamics during wakefulness, as ΔF/F□signals via iGABASnFR2, together with hippocampal local field potential (LFP) and electromyography (EMG) traces verifying wakefulness.

**Figure 1.**
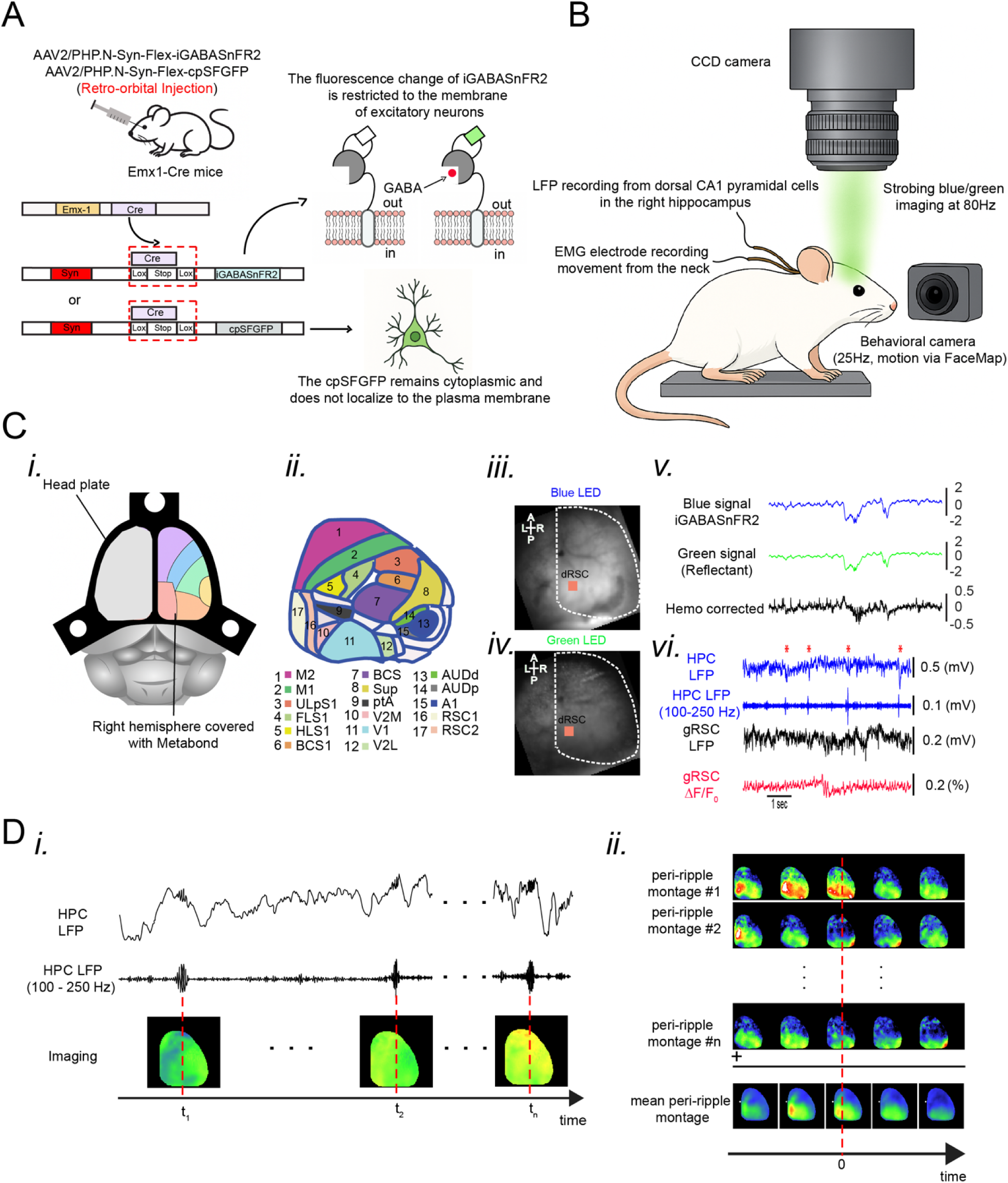
Experimental protocol for imaging neocortical GABA dynamics during sleep and hippocampal ripples. (A) Schematic of viral delivery and genetic targeting. Emx1-Cre mice received retro-orbital injections of either AAV2/PHP.eB-Syn-Flex-iGABASnFR2 or AAV2/PHP.eB-Syn-Flex-cpSFGFP as a control. In Emx1-Cre mice, Cre recombinase drives expression of the GABA sensor iGABASnFR2 or control construct specifically in cortical excitatory neurons. iGABASnFR2 is membrane-bound and reports extracellular GABA binding via fluorescence increases, whereas cpSFGFP is cytosolic and does not respond to GABA. (B) Diagram of the in vivo imaging and recording setup. A CCD camera captures widefield fluorescence from the dorsal cortex through a cranial window. Imaging is synchronized to alternating blue (470 nm) and green (530 nm) LED strobes at 80 Hz for excitation of iGABASnFR2 and collection of reflectance signals, respectively. Simultaneous recordings include hippocampal LFPs from dorsal CA1 to detect SWRs, neck EMG for movement and arousal state, and a behavioral camera (25 Hz) for motion tracking via FaceMap. (C) Imaging coverage and signal correction. (i)Schematic showing the imaging configuration: the right hemisphere was exposed for imaging, which was sealed it with Metabond, while the left hemisphere remained intact. (ii) Schematic showing the 17 identified cortical regions used in ROI-based analyses. (iii–iv) Example cortical images under blue and green LED illumination. The red box marks the dorsal retrosplenial cortex (dRSC), one of the key analysis regions. (v) Traces from the same site showing raw iGABASnFR2 fluorescence (blue), reflectance (green), and the hemodynamically corrected signal (black). (vi) Example traces showing simultaneous hippocampal LFPs (broadband and ripple-filtered), local dRSC LFP, and corrected GABA fluorescence from dRSC. Detected SWRs are aligned with both GABA and LFP signals. (D) Alignment and averaging of peri-ripple GABA activity. (i) Example SWRs from the ripple-band filtered LFP (100–250 Hz) are aligned with concurrent widefield GABA imaging frames. (ii) For each detected SWR, peri-event GABA imaging frames are extracted, aligned to the ripple center (t = 0), and assembled into individual montages. Mean peri-ripple maps are generated across events to characterize spatiotemporal GABA dynamics around SWRs.

### Cortical GABA Dynamics Across Sleep-Wake State Transitions

To examine neocortical GABA dynamics during state transitions, we used unilateral wide-field imaging of iGABASnFR2 fluorescence. Representative spatial maps and time-series traces were used to compare cortical GABA activity during NREM-to-wake and wake-to-NREM transitions.

### NREM to awake Transition

**Figure 2Ai** shows cortical GABA activity during the transition from NREM sleep to wakefulness. During this transition, z-scored ΔF/F increased across the dorsal cortex, indicating high extracellular GABA levels at wake onset. This increase is quantified in **Figure 2Aii**. Several regions, including V1, RSC2, and FLS1, showed a gradual rise in GABA activity that began a few seconds before wake onset and peaked after the transition. Coherence analysis also showed stronger inter-regional synchrony in the delta and low-theta frequency bands during this transition (**Figure S1iii**). Similarly, functional connectivity increased from more localized correlations during NREM sleep to broader cortical coupling during wakefulness (**Figure S2iv**).

**Figure 2.**
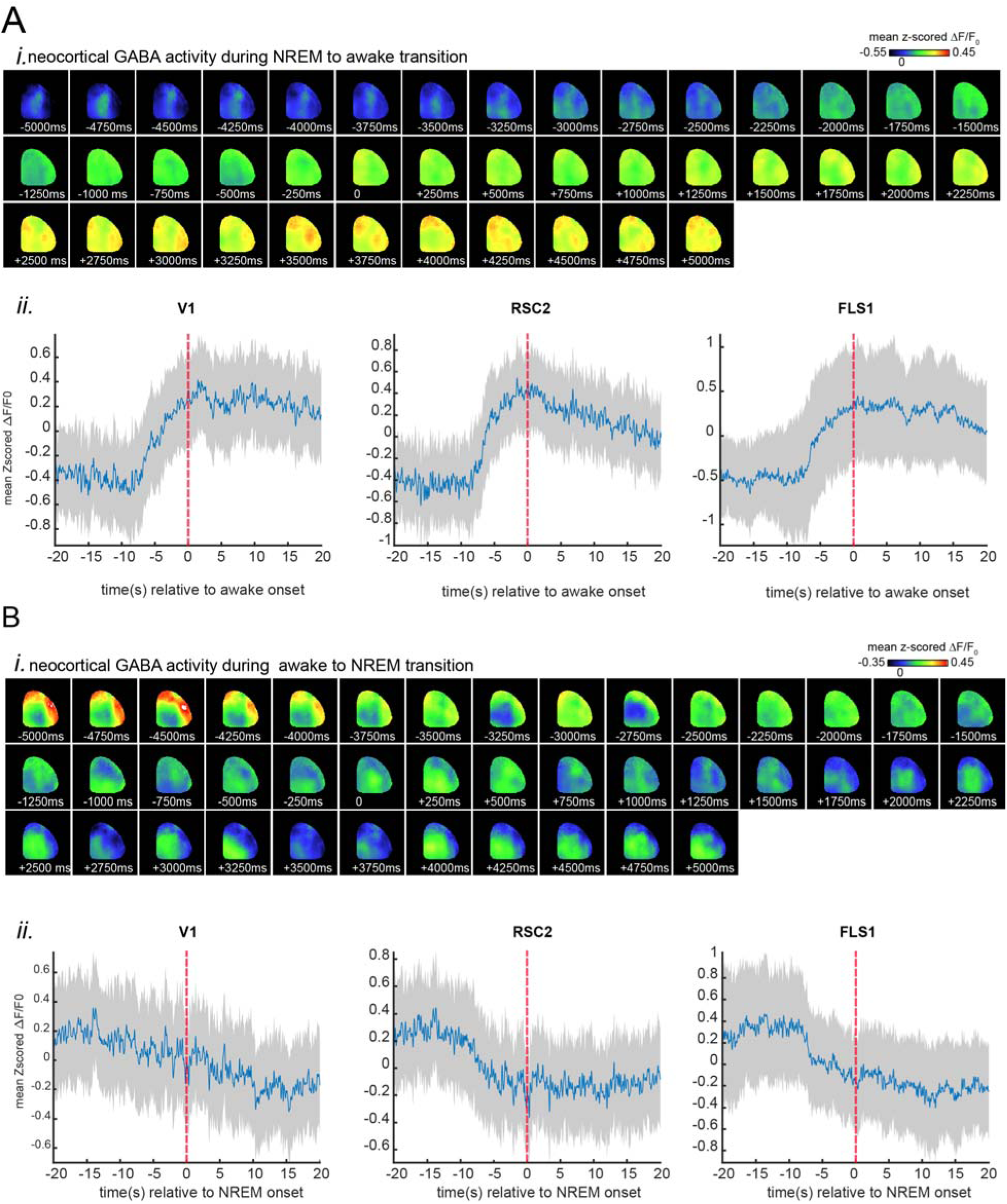
Neocortical GABA activity during NREM to Awake and Awake to NREM . (A) Transition from NREM to Awake. (i) Representative montage of mean neocortical GABA activity during wake-NREM transition recorded with the GABA-sensing fluorescent reporter iGABASnFR2. Images present 5-second time bins, and zero s-time marks wake onset. Data were scaled and z-scored to the shown color bars (n=5). (ii) Representative sample traces of iGABASnFR2 signals from regions indicated in (i). The plots present the mean optical signals recorded in 3 × 3-pixel boxes (∼0.04 mm²) throughout cortical regions. Width of shading surrounding each plot indicates standard error of the mean (SEM). The time 0 s indicates wake onset. (B) Transition from Awake to NREM. (i) Representative montage of mean neocortical GABA activity during wake-NREM transition. Images present 5-second time bins, and zero s-time marks NREM onset. Data were scaled and z-scored to the shown color bars (n=5). (ii) Representative sample traces of iGABASnFR2 signals from regions indicated in (i). The plots present the mean optical signals recorded in 3 × 3-pixel boxes (∼0.04 mm²) throughout cortical regions. Shaded areas surrounding each trace indicate reproducibility, which is confirmed. The time 0 s indicates NREM sleep onset.

### Awake to NREM Transition

**Figure 2Bi** shows cortical GABA activity during the transition from wakefulness to NREM sleep. During this transition, GABA fluorescence gradually decreased across the neocortex. This decrease is shown in **Figure 2Bii**, where V1, RSC2, and FLS1 all showed a progressive reduction in GABA activity beginning several seconds before NREM onset and continuing into early NREM sleep. Coherence analysis showed stronger low-frequency synchrony after entry into NREM sleep, consistent with the emergence of slow oscillatory network activity (**Figure S1i**). Functional connectivity also became more spatially localized during NREM compared with the broader cortical coupling seen during wakefulness (**Figure S2iii**).

### NREM to REM Transition

**Figure 3Ai** shows cortical GABA activity during the transition from NREM sleep to REM sleep. Compared with other state transitions, the GABA signal was more spatially variable, with some cortical regions showing brief increases while others remained relatively stable. In **Figure 3Aii**, V1, RSC2, and FLS1 showed a modest, transient rise in GABA activity near the transition, followed by a gradual decrease during REM sleep. Coherence analysis showed reduced inter-regional synchrony during REM compared with NREM sleep (**Figure S1iv**). Functional connectivity maps also showed weaker correlations across cortical regions during REM (**Figure S2ii**).

**Figure 3.**
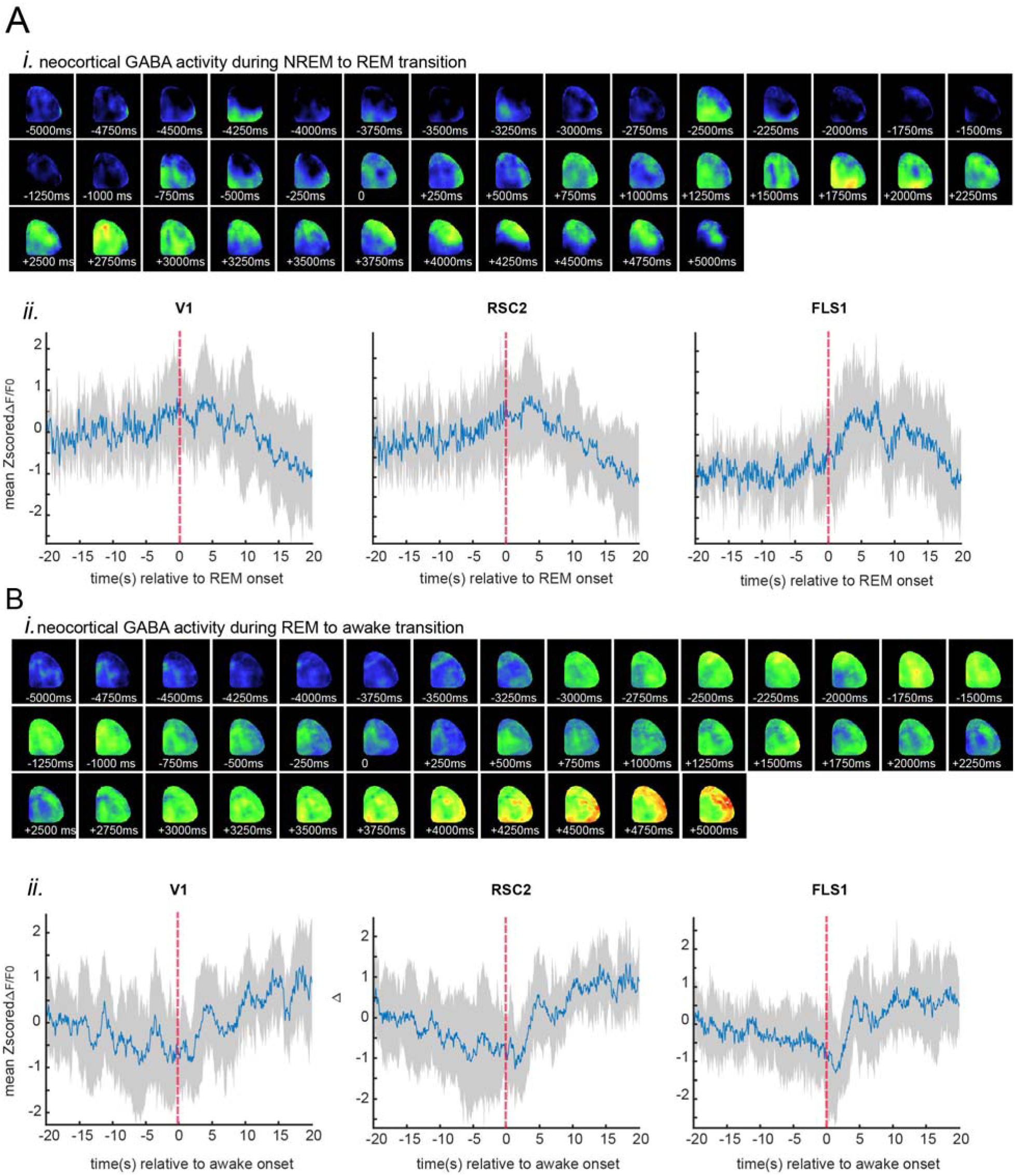
Neocortical GABA Activity During NREM to REM and REM to Awake Transitions. (A) Transition to REM. (i) Montage of mean neocortical GABA activity recorded using the GABA-sensing fluorescent reporter iGABASnFR2 during natural sleep. Images represent 5-second epochs, and zero s-time represents REM onset. Data were z-scored and normalized to the color bars shown (n=5). (ii) Representative traces of iGABASnFR2 signals from regions marked in (i) in an exemplary animal. The plots show the mean optical signals recorded in 3 × 3-pixel boxes (∼0.04 mm²) within the visual (V1), retrosplenial (RSC), and forelimb somatosensory (FLS1; magenta) cortices. The thickness of the shading around each plot gives the SEM. The time 0 s represents REM onset. (B) Transition to wake at wake-onset. (i) Montage of mean neocortical GABA activity recorded using the GABA-sensing fluorescent reporter iGABASnFR2 during wake-onset transition from REM. Images represent 5-second epochs, and zero s-time represents wake-onset. Data were z-scored and normalized to the color bars shown (n=5). (ii) Representative traces of iGABASnFR2 signals from regions marked in (i) in an exemplary animal. The plots show the mean optical signals recorded in 3 × 3-pixel boxes (∼0.04 mm²) within the visual (V1), retrosplenial (RSC2), and forelimb somatosensory (FLS1; magenta) cortices. The thickness of the shading around each plot gives the SEM. The time 0 s represents wake-onset.

### REM to Wake Transition

**Figure 3Bi** shows cortical GABA activity during the transition from REM sleep to wakefulness. At the transition, the GABA signal briefly decreased and then increased again. This transient decrease is shown in **Figure 3Bii** and was most visible in V1 and FLS1. This pattern suggests a short period of reduced cortical GABA signaling as the brain exits REM sleep. Coherence analysis showed increased inter-regional synchrony in the delta and low-theta frequency bands during this transition (**Figure S1ii**), consistent with a shift toward wake-like network activity. Functional connectivity analysis also showed stronger inter-regional correlations after REM, especially within anterior sensory-motor cortical regions (**Figure S2i**).

### State-dependent cortical GABA dynamics around sharp-wave ripples

Previous studies have shown that hippocampal sharp-wave ripples (SWRs) coordinate hippocampal–cortical communication during offline brain states, such as sleep and quiet wakefulness ^43,45–48^. It has been shown that the disruption of SWRs impairs memory consolidation, indicating their importance for brain function ^49–50^. These events provide a useful model for examining how cortical GABA activity is modulated in a localized and time-specific manner. Using the iGABASnFR2 sensor, we combined unilateral wide-field imaging of neocortical extracellular GABA with CA1 local field potential recordings. This allowed us to examine the spatial and temporal patterns of cortical GABA dynamics around hippocampal SWRs during natural sleep and wakefulness. **Figure 4A** shows the spatial and temporal patterns of neocortical GABA activity around SWRs across different brain states. During NREM sleep, GABA activity began to increase about 100 ms before the SWR center (t = 0; **Figure 4Ai**; Video 7). This activation was strongest in the retrosplenial cortex (RSC) and peaked before the SWR center (Figure 4Bi). The response showed a medial-to-lateral pattern, with medial regions such as RSC activating earlier and more strongly. In contrast, lateral cortical areas showed reduced GABA activity before the SWR center. During wakefulness, cortical GABA activity showed a weak increase before the SWR center and continued to rise after t = 0 (**Figure 4Aii**). The response reached its maximum about 450–500 ms after the SWR (**Figure 4Biii–iv**). Spatially, the activity progressed from lateral regions, including visual and somatosensory cortex, toward medial cortical areas. Compared with NREM sleep, pre-ripple decreases were weaker and less widespread, indicating that SWR-related GABA modulation differs across brain states.

**Figure 4.**
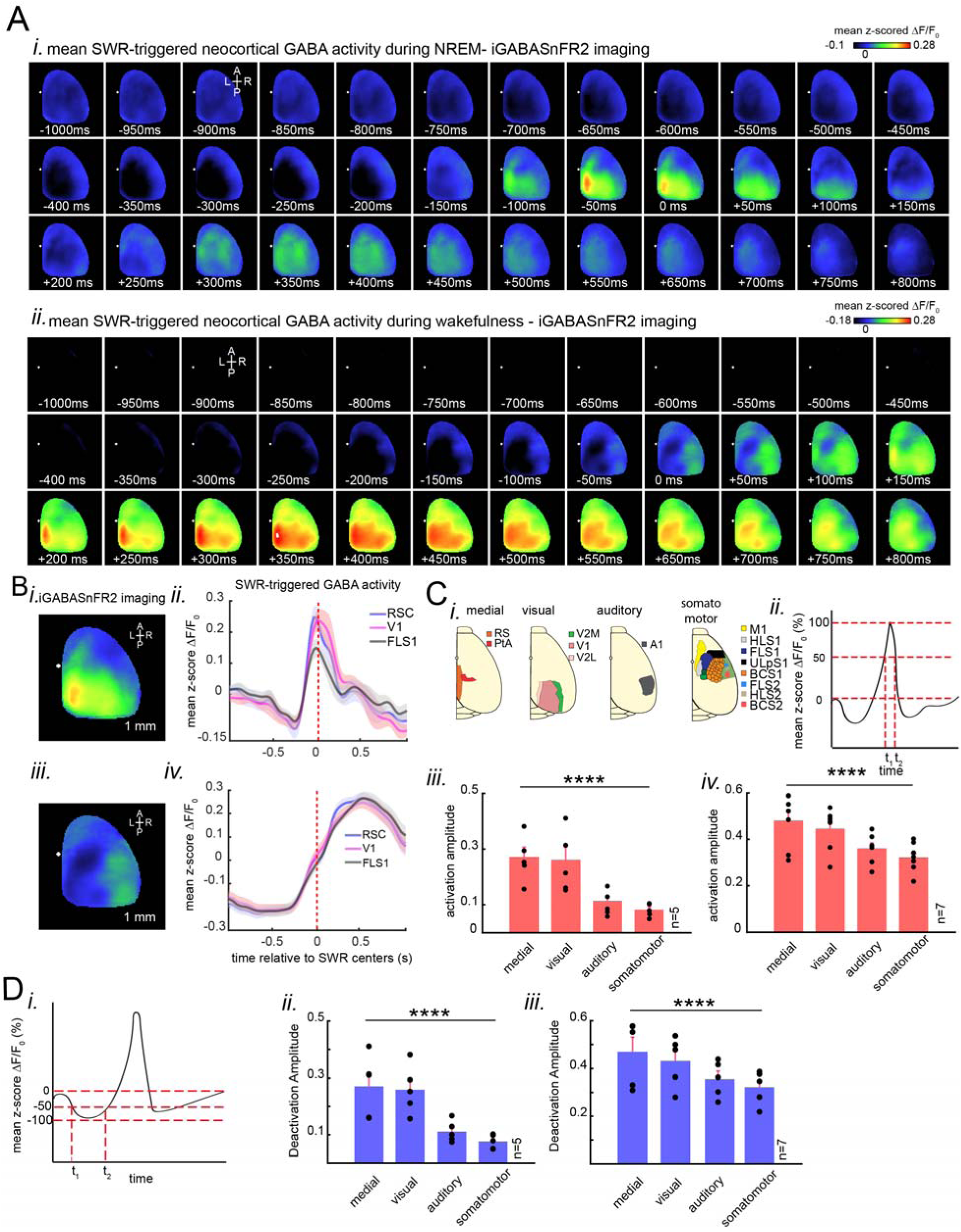
Peri-SWR neocortical GABA activity: activation and deactivation patterns across sleep and wake states. (A) Characteristic montage of mean peri-SWR neocortical GABA activity, as visualized with the GABA-sensing fluorescent reporter iGABASnFR2, during head-restrained natural sleep (i) and wakefulness (ii). Time 0 marks SWR centers. Images z-scored and scaled to respective color bars. (B) (i–iv) Sample traces of iGABASnFR2 signals. Panels (Bi, Bii) show natural sleep traces, and (Biii, Biv) show wake traces, taken from sample regions in panels (Ai) and (Aii). Traces show mean optical signals from 3 x 3-pixel boxes (∼0.04 mm^2^) within retrosplenial (blue), visual (black), and forelimb somatosensory (magenta) cortices. Shaded areas around regions of each plot are SEM. SWR centers mark time 0. (C) (i) Four large neocortical subnetworks (medial, visual, auditory, and somatomotor), structurally delimited. (ii) Quantification of activation amplitude schema, defined as mean within-whole-width-at-half maximum (t1 to t2) of the mean. (iii–iv) Grand average of activation amplitudes across subnetworks during sleep (n = 7 animals) and wake (n = 7 animals) ordered in decreasing order. Each data point represents average activation amplitude for all regions within a subnetwork for one animal. Statistical Analysis: Sleep data also revealed a significant subnetwork effect (F (3,12) = 21.608, p = 3.96 × 10□□, Greenhouse-Geisser corrected p = 0.0017), with posthoc differences between auditory and medial (p = 0.0305), auditory and visual (p = 0.0146), and medial and somatomotor (p = 0.0366). Awake data had a significant subnetwork influence (F(3,18) = 30.051, p = 3.18 × 10□□, Greenhouse-Geisser adjusted p = 7.03 × 10□□), and post-hoc differences were found between auditory and medial (p = 0.0075), auditory and visual (p = 0.0025), medial and somatomotor (p = 0.0027), and somatomotor and visual (p = 0.0053). (D) (i–iii) neocortical deactivations preceding SWRs. (Dii, Diii) show results for sleep and wakefulness respectively. Deactivation amplitudes were rectified for group comparison, with higher values indicating more substantial deactivation. Bar plots depict mean ± SEM. Statistical Analysis: Sleep data recorded a significant subnetwork effect (F (3,12) = 21.608, p = 3.96 × 10□□, Greenhouse-Geisser corrected p = 0.0017), and posthoc differences between auditory and visual (p = 0.015), auditory and medial (p = 0.031), and somatomotor and visual (p = 0.010) regions. There are five supplementary panels in the figure. Awake data recorded a significant subnetwork effect (F (3,12) = 13.869, p = 0.00033, Greenhouse-Geisser corrected p = 0.0047), and posthoc differences between auditory and visual (p = 0.032), medial and somatomotor (p = 0.041), and visual and somatomotor (p = 0.061) regions.

To further examine the spatial organization of peri-SWR GABA activity, we analyzed four major neocortical subnetworks: medial, visual, auditory, and somatomotor regions, as shown in **Figure 4Ci**. Activation amplitudes were determined by taking the mean signal across the full width at half-maximum (from t□ to t□), as in **Figure 4Cii**. Repeated-measures ANOVA showed that peri-SWR GABA activity differed across cortical subnetworks during both wakefulness and NREM sleep. During wakefulness, peri-SWR GABA activity differed significantly across cortical subnetworks (**Figure 4Civ**; repeated-measures ANOVA: F (3,18) = 30.051, p = 3.18 × 10□□; Greenhouse–Geisser adjusted p = 7.03 × 10□□). Post-hoc analyses showed that the medial subnetwork had significantly stronger activation than the auditory (p = 0.0075) and somatomotor (p = 0.0027) subnetworks. The visual subnetwork also showed significantly stronger activation than the auditory (p = 0.0025) and somatomotor (p = 0.0053) subnetworks. During NREM sleep, peri-SWR GABA activity also differed significantly across cortical subnetworks (**Figure 4Cv**; repeated-measures ANOVA: F(3,12) = 21.608, p = 3.96 × 10□□; Greenhouse–Geisser adjusted p = 0.0017). Post-hoc analyses showed that the medial subnetwork had significantly stronger activation than the auditory (p = 0.0305) and somatomotor (p = 0.0366) subnetworks. The visual subnetwork showed intermediate activation but was still significantly higher than the auditory (p = 0.0146) and somatomotor (p = 0.0366) subnetworks. Deactivation amplitudes also showed state-specific patterns during wakefulness and NREM sleep (**Figure 4D**). During wakefulness, deactivation amplitudes were relatively small but still differed significantly across subnetworks (**Figure 4Di**; repeated-measures ANOVA: F (3,12) = 13.869, p = 0.00033; Greenhouse–Geisser adjusted p = 0.0047). Post-hoc tests showed that the auditory subnetwork had significantly less deactivation than the visual (p = 0.032) and medial (p = 0.041) subnetworks. The visual subnetwork also showed greater deactivation than the somatomotor subnetwork, although this comparison did not reach significance (p = 0.061). During NREM sleep, deactivation amplitudes were larger and differed significantly across subnetworks (**Figure 4Dii**; repeated-measures ANOVA: F (3,12) = 21.608, p = 3.96 × 10□□; Greenhouse–Geisser corrected p = 0.0017). Post-hoc tests showed significant differences between the auditory and visual subnetworks (p = 0.015), auditory and medial subnetworks (p = 0.031), and somatomotor and visual subnetworks (p = 0.010).

To further examine state-dependent modulation, we measured GABA activation and deactivation amplitudes across neocortical regions during wakefulness and NREM sleep (**Figure S3A**). Activation and deactivation patterns were broadly similar across states, as supported by correlation analyses (**Figure S3B**). At the subnetwork level, activation and deactivation amplitudes did not differ significantly between NREM sleep and wakefulness, indicating that subnetwork-specific patterns of GABA modulation were largely preserved across these states (**Figure S3C**). For activation amplitudes, mixed repeated-measures ANOVA showed a significant main effect of state, with higher overall activation during NREM sleep than wakefulness (F (1,10) = 28.266, p = 0.00034). There was also a significant main effect of subnetwork (F (3,30) = 51.895, p = 5.50 × 10□¹²), indicating that activation amplitudes differed across cortical regions. However, the state × subnetwork interaction was not significant (F (3,30) = 1.605, p = 0.209), suggesting that the relative subnetwork pattern was preserved across states. For deactivation amplitudes, mixed repeated-measures ANOVA also showed a significant main effect of state, with stronger deactivation during NREM sleep than wakefulness (F (1,8) = 18.571, p = 0.0026). The main effect of subnetwork was significant as well (F (3,24) = 35.134, p = 6.08 × 10□□), showing that deactivation amplitudes differed across cortical subnetworks. The state × subnetwork interaction was not significant (F (3,24) = 1.686, p = 0.197), indicating that subnetwork-specific deactivation patterns remained consistent across brain states.

### SWR-associated cortical GABA activity propagates differently across brain states

We next examined when peak cortical GABA activation occurred across different cortical areas. Peak timing (t□) was defined as the time of maximum GABA signal relative to the SWR center (t = 0; **Figure 5Ai**). This analysis allowed us to map the spatiotemporal organization of SWR-related GABA dynamics during NREM sleep and wakefulness.

**Figure 5.**
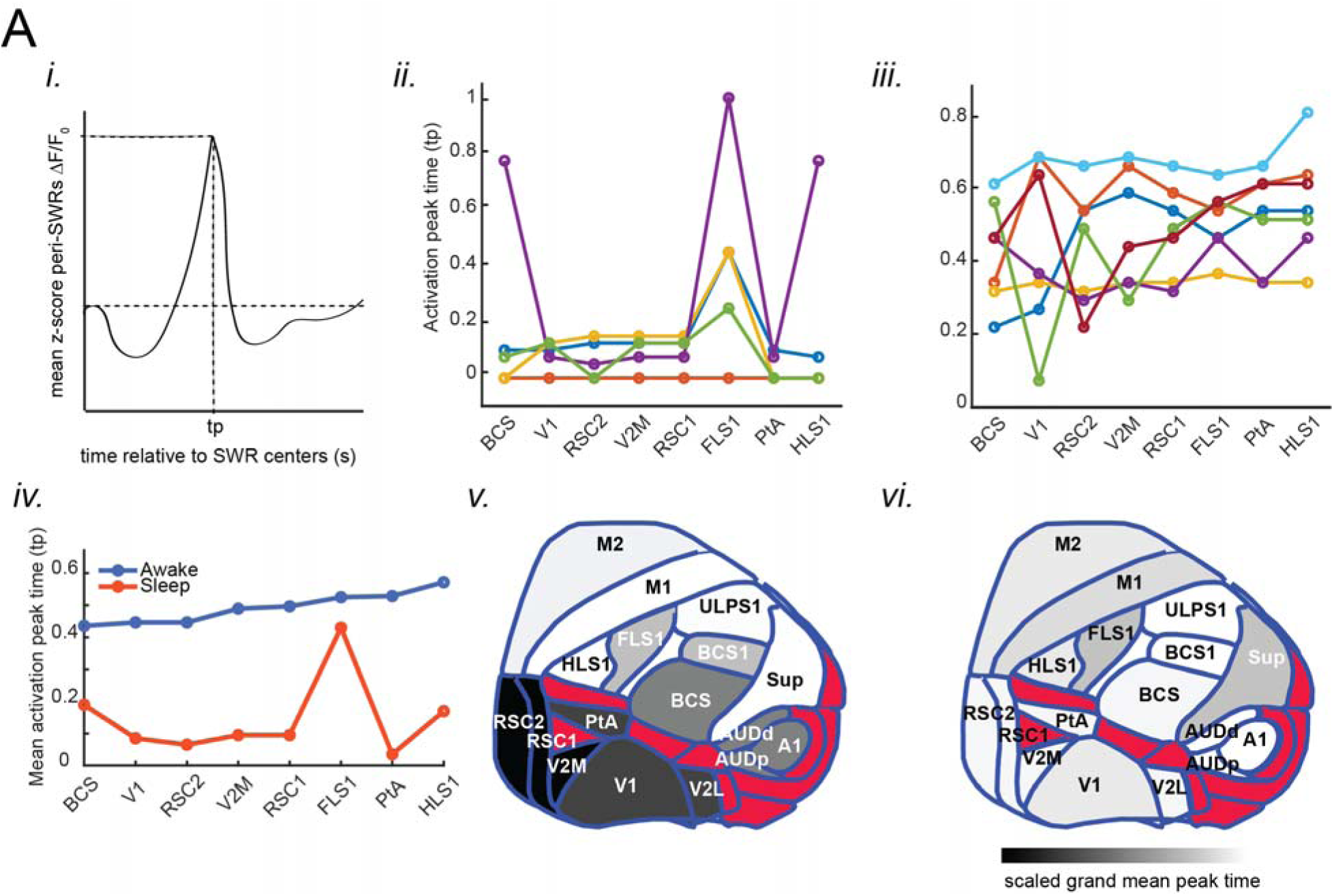
Cortical regions exhibit state-dependent peak GABA activation dynamics around SWRs during natural sleep and wakefulness. (A) (i) Peak GABA activation time (tp) for peri-SWR in each cortical region was determined as peak time of mean peri-SWR GABA activity trace (blue trace) and provides an index of peak GABA activation time relative to center of SWR. (ii) Sleep Condition: Repeated-measures ANOVA revealed significant regional variation in tp (F(10,40) =5.0973, p=8.9111e−05). A statistically significant positive linear trend (slope=0.011273□s/region,p=0.047249) discloses medial-to-lateral progression of tp during sleep, consistent with a subtle medial-to-lateral propagation of GABA activation. (iii) Awake Condition: Repeated-measures ANOVA revealed strong and statistically significant regional variation in tp (F(10,60) =20.156, p=1.116e−15). Post-hoc linear trend analysis confirmed a statistically significant positive slope (slope=0.0272□s/region, p=0.024326), which discloses a consistent and systemic delayed, tp from medial toward lateral regions in wake stage. (iv) Post-hoc Linear Trend Analysis: Linear trend analysis confirmed state-dependent dynamics. In wake, a statistically significant positive slope (0.02724□s/region=0.024326) discloses a robust medial-to-lateral gradient. In sleep, a negligibly but statistically significant smaller slope (0.011273□s/region=0.047249) reveals less strenuous regional variations. The corresponding graph reveals the result, which shows steep and consistent regional variation in awake data compared with sleep. (v) Spatial maps during sleep (n=5) reveal higher earlier peak tp in medial regions like RSC1, RSC2, and PtA, and subsequent lateral propagation of activation. The result discloses an orchestrated medial-to-lateral propagation of GABAergic activation during natural sleep. (vi) Spatial maps for wakefulness (n=7) indicate delayed activation in medial regions and a more homogeneous distribution of tp values throughout cortical regions. The pattern indicates characteristic cortical dynamics in wakefulness, with a strong medial-to-lateral gradient in GABA activation timing.

During NREM sleep, peak timing varied across cortical regions (**Figure 5Aii**), as confirmed by repeated-measures ANOVA (F (10,40) = 5.0973, p = 8.91 × 10□□). Linear trend analysis showed a weak but significant medial-to-lateral gradient in activation timing (slope = 0.0113 s/region, p = 0.047). Spatial maps supported this pattern, with medial regions such as RSC1 and RSC2 reaching peak activation earlier than lateral cortical areas (**Figure 5Av**). During wakefulness, peak timing showed stronger and more organized regional variation (**Figure 5Aiii**), as confirmed by repeated-measures ANOVA (F (10,60) = 20.156, p = 1.12 × 10□¹□). Linear trend analysis showed a steeper gradient than in NREM sleep (slope = 0.0272 s/region, p = 0.024). Spatial maps showed an inverted pattern, with medial regions reaching peak GABA activation later than lateral areas (**Figure 5Avi**). Direct comparison of slope values confirmed that this timing gradient was stronger during wakefulness than during NREM sleep (**Figure 5Aiv**).

Overall, these findings show that peak cortical GABA timing differs by brain state, with a gradual pattern during NREM sleep and a more structured pattern during wakefulness. Similar rise times across regions and states suggest that these differences mainly reflect when activation begins or peaks, rather than how fast the GABA signal rises (**Figure S4A**). During wakefulness, peak timing was positively correlated with lateral cortical position (r = 0.558, p = 0.011; **Figure S4Bii**), consistent with a lateral-to-medial progression of GABA activation. In NREM sleep, peak timing was negatively correlated with cortical position (r = −0.391, p = 0.040; **Figure S4Biii**), indicating earlier activation in medial regions.

### Spatial and temporal modes of peri-ripple cortical GABA activity

To examine how hippocampal ripples are associated with cortical GABA dynamics, we decomposed peri-ripple iGABASnFR2 signals into spatial and temporal modes using singular value decomposition (SVD; **Figure 6A, B**). Each component represents a pattern of cortical regions that change together over time relative to the ripple center (0 s). In both wakefulness and NREM sleep, the first component showed a broad, near-global cortical pattern with a gradual increase after the ripple. Components 2–5 captured more localized, region-specific patterns that differed between brain states. During NREM sleep, spatial maps showed broad cortical engagement, with stronger involvement of medial regions such as the retrosplenial cortex (**Figure 6A**). The temporal traces showed slow, sustained increases around ripple time, suggesting a more widespread and prolonged GABA modulation. During wakefulness, spatial maps emphasized lateral sensory and motor regions (**Figure 6B**). The temporal modes showed smaller biphasic changes, indicating faster and more localized peri-ripple GABA modulation. To link these components to regional activity, we reconstructed peri-ripple GABA traces from five cortical areas: RSC, barrel cortex (BC), primary visual cortex (V1), auditory cortex (AUD), and forelimb somatosensory cortex (FLS1) (**Figure 6C, D**). During wakefulness, BC and V1 showed clear GABA transients with slightly higher amplitudes than medial regions. During NREM sleep, RSC showed the most prolonged and elevated response. Together, these results show that hippocampal SWRs are associated with a broad increase in cortical GABA activity, but the spatial and temporal pattern depends on brain state. During wakefulness, GABA modulation was faster and more localized, mainly involving lateral sensory regions. During NREM sleep, GABA activation was slower and more widespread, with stronger involvement of medial regions such as RSC.

**Figure 6.**
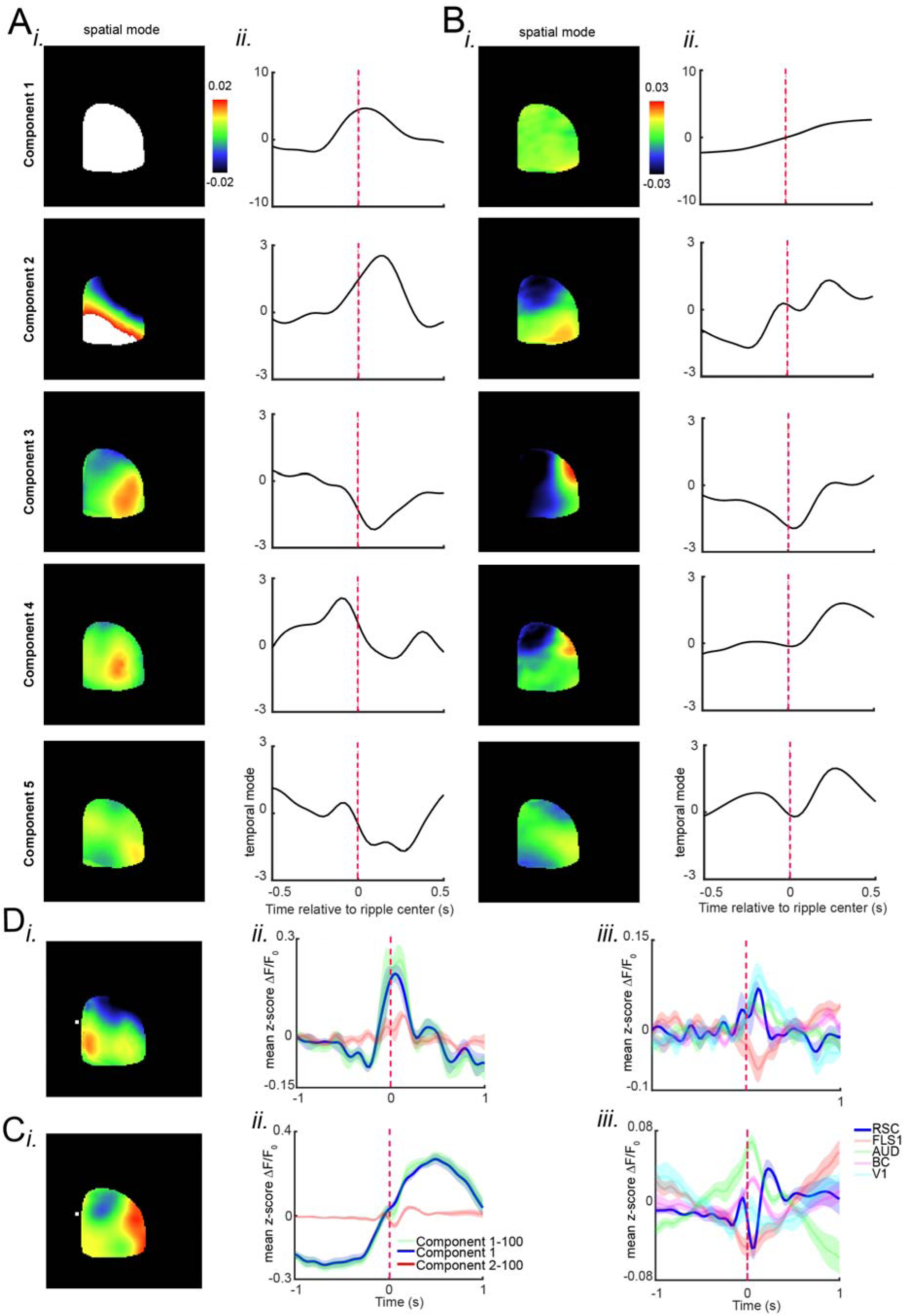
Spatial and Temporal Modes of Peri-Ripple iGABASnFR2 Activity and Region-Specific Dynamics. (A, B) Spatial (i) and temporal (ii) modes of the five largest singular components of the peri-ripple combined iGABASnFR2 activity stack in **sleep (A)** and **wakeful (B)** epochs. The spatial mode of the first component exhibits an absence of sharp topographical features, consistent with a widespread increase of the iGABASnFR2 signal following the ripple, which dominates the overall GABA signal and corresponding temporal mode. This large-scale response is invariant across states, with the temporal amplitude of the first component being one order of magnitude larger than the others, thus revealing its predominance in describing the dynamics of global peri-ripple GABA activity. Components 2–5 demonstrate **state-dependent differences** in the spatiotemporal structure of GABA activity. **During sleep (A)**, spatial modes of Components 2–5 reveal less spatially localized patterns with increased associative area activity (e.g., retrosplenial cortex, RSC) and decreased sensory area activity. Corresponding temporal profiles during sleep show slower dynamics without sharp peaks. **During wakefulness (B)**, spatial maps of Components 2–5 illustrate localized GABA activity, with particularly strong activity in sensory areas such as the barrel cortex (BC) and visual cortex (V1). Their temporal profiles reveal rapid, sharp peaks at fixed times of ripple initiation, indicating specific and region-dependent GABAergic control. (C, D) Peri-ripple regional GABA activity reconstructed for significant regions of interest (ROIs)—the retrosplenial cortex (RSC), barrel cortex (BC), visual cortex (V1), auditory cortex (AUD), and forelimb somatosensory cortex (FLS1)—reveal **state-dependent peri-ripple dynamics**. **During sleep (C)**, RSC shows prolonged and sizable GABA activity relative to sensory areas. **During wakefulness (D)**, BC and V1 reveal well-defined and high-amplitude peaks in their temporal profiles (Dii), and statistical comparison (Diii) shows significantly increased sensory responses compared to RSC. Shading in temporal profiles (Cii, Dii) denotes ±SEM. Statistical comparisons emphasize the relative enhancement of RSC activity during sleep and the sensory-dominant responses during wakefulness.

## Discussion

### Summary of the Study

In this study, we examined how neocortical GABA dynamics are spatially and temporally organized around internally generated hippocampal sharp-wave ripples (SWRs) across behavioral states. Using wide field iGABASnFR2 imaging together with simultaneous CA1 LFP recordings, we monitored extracellular GABA activity across the dorsal cortex during natural sleep and wakefulness. We first characterized cortical GABA dynamics during transitions between wakefulness, NREM sleep, and REM sleep. We then examined peri-SWR GABA activity during NREM sleep and wakefulness. Our results showed that SWRs are associated with structured cortical GABA modulation, including broad activation, regional deactivation, and state-dependent propagation patterns. Specifically, GABA activity showed a medial-to-lateral organization during NREM sleep and a lateral-to-medial organization during wakefulness.

### Cortical GABA Dynamics During State transitions

First, we examined the changes in cortical GABA levels that take place during spontaneous behavior state transitions, and we complemented this analysis with coherence and correlation-based network methods to reveal their implications at the circuit level. As the brain transitioned from NREM sleep to wakefulness, we observed a dramatic increase in GABA activity, which was replicated across different cortical areas (Figure 2Ai–ii), and increased inter-areal coherence and functional connectivity (Figures S1iii, S2iv). This increase in inhibitory tone, coupled with increased synchrony, can reflect an active gating or stabilization process during the arousal process, promoting the coordination of sensory and motor pathways as the brain prepares for the state of behavioral responsiveness^52–54^. In contrast, transitions from wakefulness to NREM sleep were correlated with a decrease in extracellular GABA (Figure 2Bi–ii), together with increases in slow frequency coherence (Figure S1i) and more tightly localized functional correlations (Figure S2iii). These are likely to represent a switch from fast, phasic inhibitory mechanisms to slower, synchronous, oscillatory modes typical of NREM sleep^55,56^, in favor of the commencement of cortical downscaling and memory consolidation. The dynamics of GABA in the neocortex across transitions involving REM sleep showed increasingly complex and varied dynamics. The transition from NREM to REM sleep was marked by modest, regionally specific increases in GABA levels, which were then followed by suppression (Figure 3Ai–ii). This transition also showed a decrease in coherence and weakening of inter-areal correlations (Figures S1iv, S2ii), consistent with the decoupled cortical dynamics characteristic of REM sleep. In contrast, the transition from REM sleep to wakefulness showed a large drop in GABA at the point of awakening (Figure 3Bi–ii), followed by rapid re-engagement, reflecting a brief release from inhibitory control ^57–60^. This transition was accompanied by a large increase in coherence and correlation strength (Figures S1ii, S2i), marking a reorganization of network synchrony required for cognitive and sensory engagement in wakefulness. Together, these findings highlight the dynamic and state-specific regulation of GABAergic tone across the neocortex, indicating that changes in inhibitory signaling are closely intertwined with macroscale network reconfiguration across sleep-wake transitions.

### Ripple-triggered GABA responses are brain-state dependent

Past work using iGluSnFR has shown that hippocampal SWRs are coordinated with state-dependent cortical excitation ^21–23^. Here we extend this by showing that extracellular GABA dynamics are also state-dependent around SWRs. During NREM sleep, medial cortical areas exhibited a brief decrease in extracellular GABA immediately before SWR onset, followed by a robust increase after the ripple, which began medially and propagated toward lateral cortex. This biphasic pattern aligns with reports that SWRs recruit retrosplenial neurons during slow-wave sleep^61^and with prior evidence for retrosplenial–hippocampal coordination in memory-related processing^62,63^. In wakefulness, extracellular GABA levels increased predominantly in lateral sensory cortices after ripple onset, whereas medial regions showed delayed or attenuated changes. Together, these results demonstrate that SWRs are accompanied by state-dependent, spatially organized changes in extracellular GABA, revealing distinct network-level coordination patterns during sleep and wakefulness. Consistent with this interpretation, the quantitative activation and deactivation analyses (Figure 4C–D) confirm that all cortical subnetworks displayed a pre-SWR decrease (deactivation) followed by a post-SWR increase (activation) in extracellular GABA, with the strongest and earliest responses in medial areas during NREM sleep and a reversed lateral-to-medial organization during wakefulness.

### Temporal gradients reveal propagation of inhibitory signals

Time-to-peak (t□) of neocortical GABA dynamics throughout different cortical areas showed medial-to-lateral propagation during NREM sleep, consistent with glutamate dynamics 19 and memory-related information flow direction between default-mode network structures. During wakefulness, direction of propagation was reversed, and propagation went from lateral to medial cortical compartments. These results are in concert with the suggestion that cortical inhibition characteristically follows excitation with a characteristic temporal delay ^1,64,65^. Neocortical inhibition in this case can function as a state-dependent gate of the hippocampal output, orchestrating temporally the inhibition control with the current sensory or cognitive demands. Directional asymmetry of propagation patterns uncovers the cortex to rescale not only the strength but also the spatiotemporal organization of inhibition in accordance with internal state.

### SVD reveals global and local components of peri-SWR GABA dynamics

SVD analysis of neocortical iGABASnFR2 signals revealed distinct spatial and temporal patterns of peri-SWR GABA dynamics. The dominant component showed a broad, near-global increase in cortical GABA activity, consistent with a coordinated network-level response to hippocampal SWRs. Higher-order components captured more localized and state-dependent patterns. During wakefulness, these components reflected brief GABA transients in lateral sensory and motor regions, suggesting more localized cortical modulation around SWRs. During NREM sleep, GABA activity was broader and more sustained, with stronger involvement of medial regions such as the retrosplenial cortex, a hippocampal-connected area involved in systems-level memory processing.

## Conclusion

In this study, mesoscale iGABASnFR2 imaging was used to map extracellular GABA dynamics across the dorsal cortex during sleep–wake transitions and around hippocampal sharp-wave ripples. Our results show that cortical GABA signaling is spatially patterned, strongly state dependent, and temporally organized around hippocampal SWRs. These findings suggest that neocortical GABA dynamics may contribute to the filtering of hippocampal output during sleep and wakefulness. Although wide field iGABASnFR2 imaging provides broad cortical coverage with high temporal resolution, it does not identify the specific interneuron classes or cortical layers responsible for the observed signals. Therefore, this approach cannot distinguish the contributions of PV+, SST+, or VIP+ interneurons, nor can it resolve deeper subcortical sources of GABAergic modulation. Future studies combining iGABASnFR2 with optogenetic manipulation, cell-type-specific labeling, or multiphoton imaging will be important for defining the circuit mechanisms underlying state-dependent and SWR-related GABA dynamics. In addition, combining GABA sensors with glutamate sensors could allow direct investigation of excitation–inhibition balance in vivo, both under normal conditions and in disease models such as epilepsy or schizophrenia.

## Acknowledgements

We thank Di Shao and the Animal Welfare Committee at the University of Lethbridge for their support in enabling the animal experiments.

## Funding

This work was supported by Alberta Innovates and the Natural Sciences and Engineering Research Council of Canada (Grant No. 390930), awarded to Majid H. Mohajerani.

## Author contributions

E.R. conceptualized the study, designed and performed the experiments, analyzed the data, and wrote the manuscript. S.T. contributed to data analysis. M.N. contributed to the investigation. B.L.M. contributed to conceptualization, supervision. M.H.M. contributed to conceptualization, resources, supervision, and funding acquisition. All authors reviewed, commented on, and approved the final manuscript.

## Competing interests

The authors declare no competing interests.

## Supplementary material

Supplementary material and methods are available upon request.

---

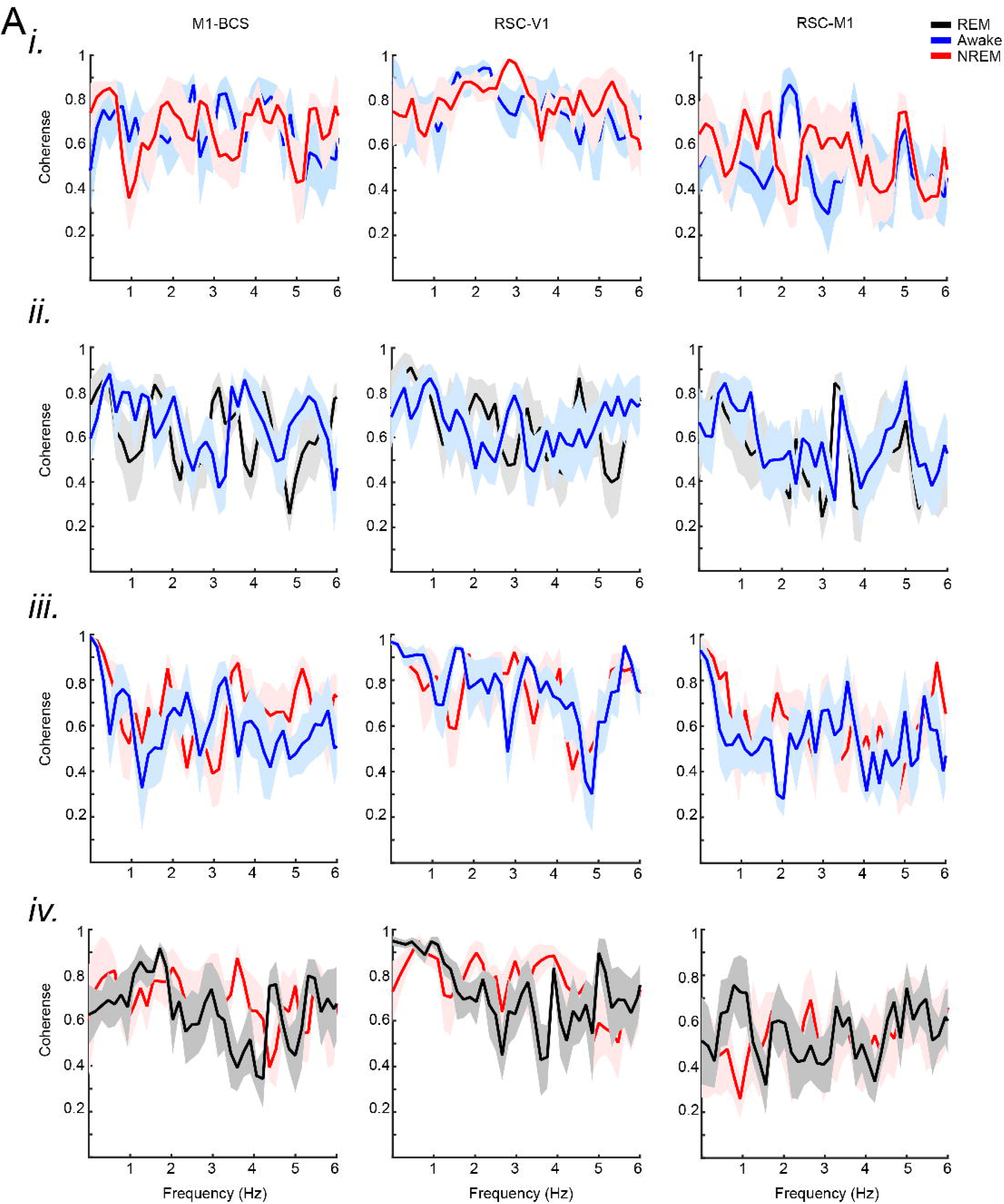

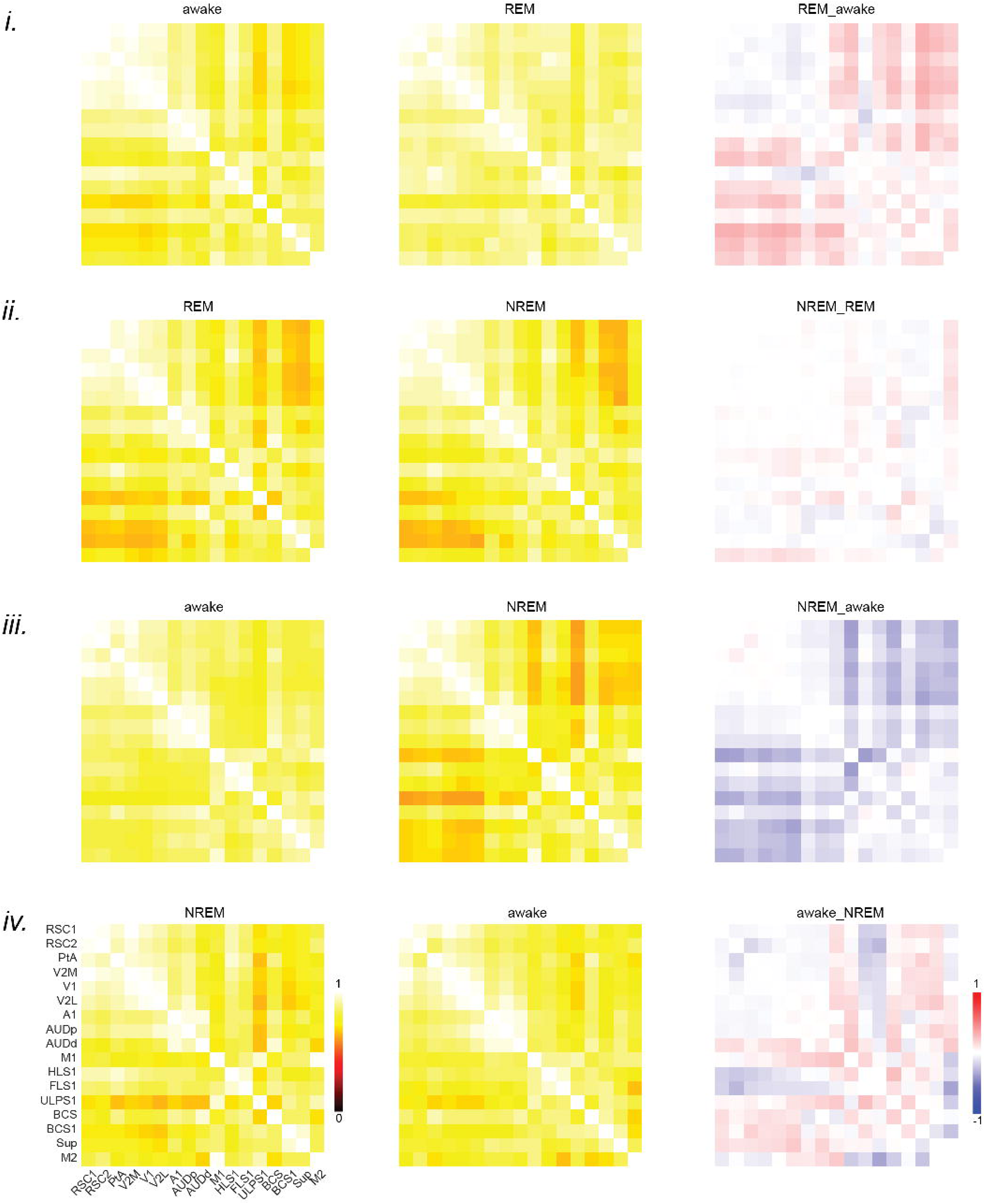

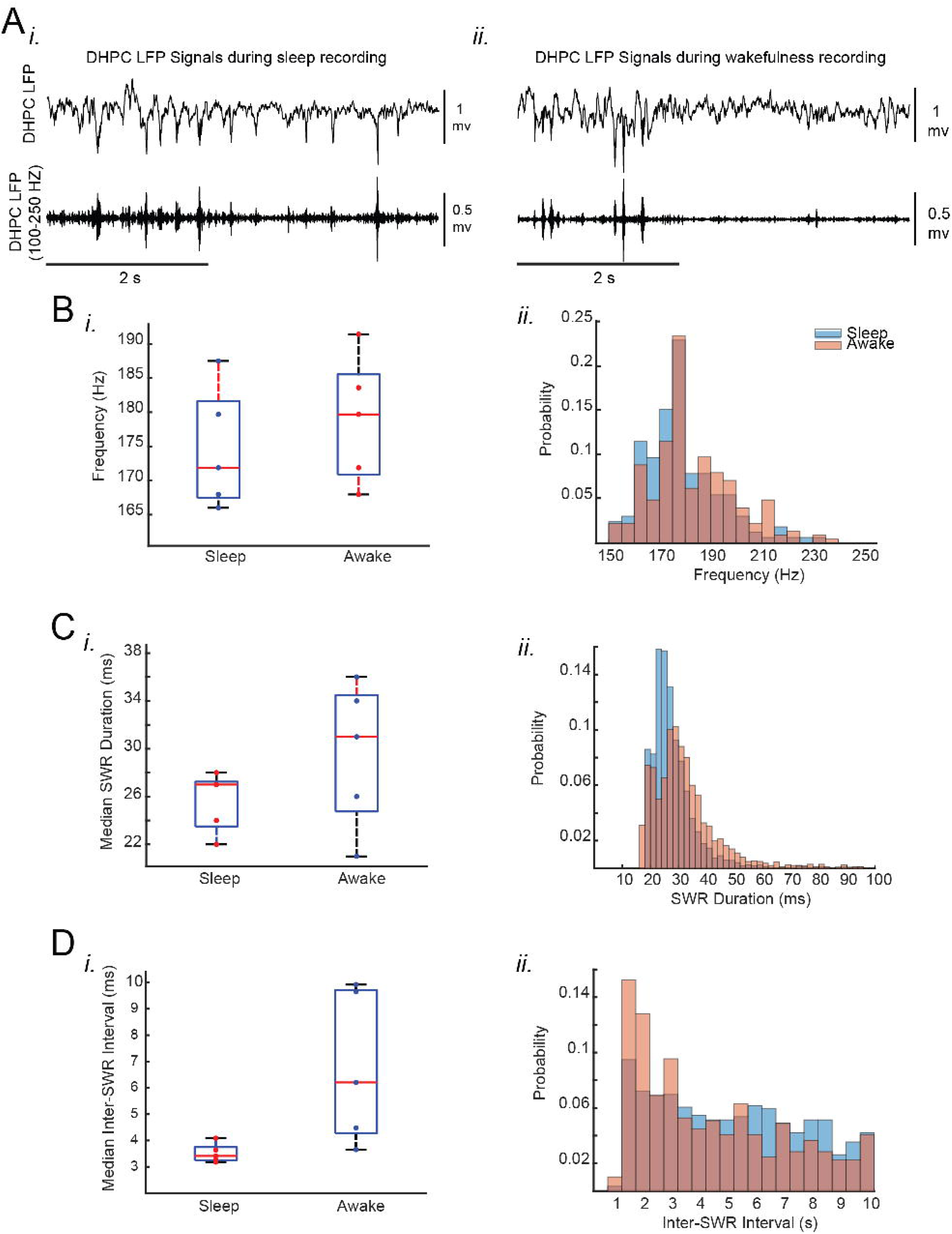

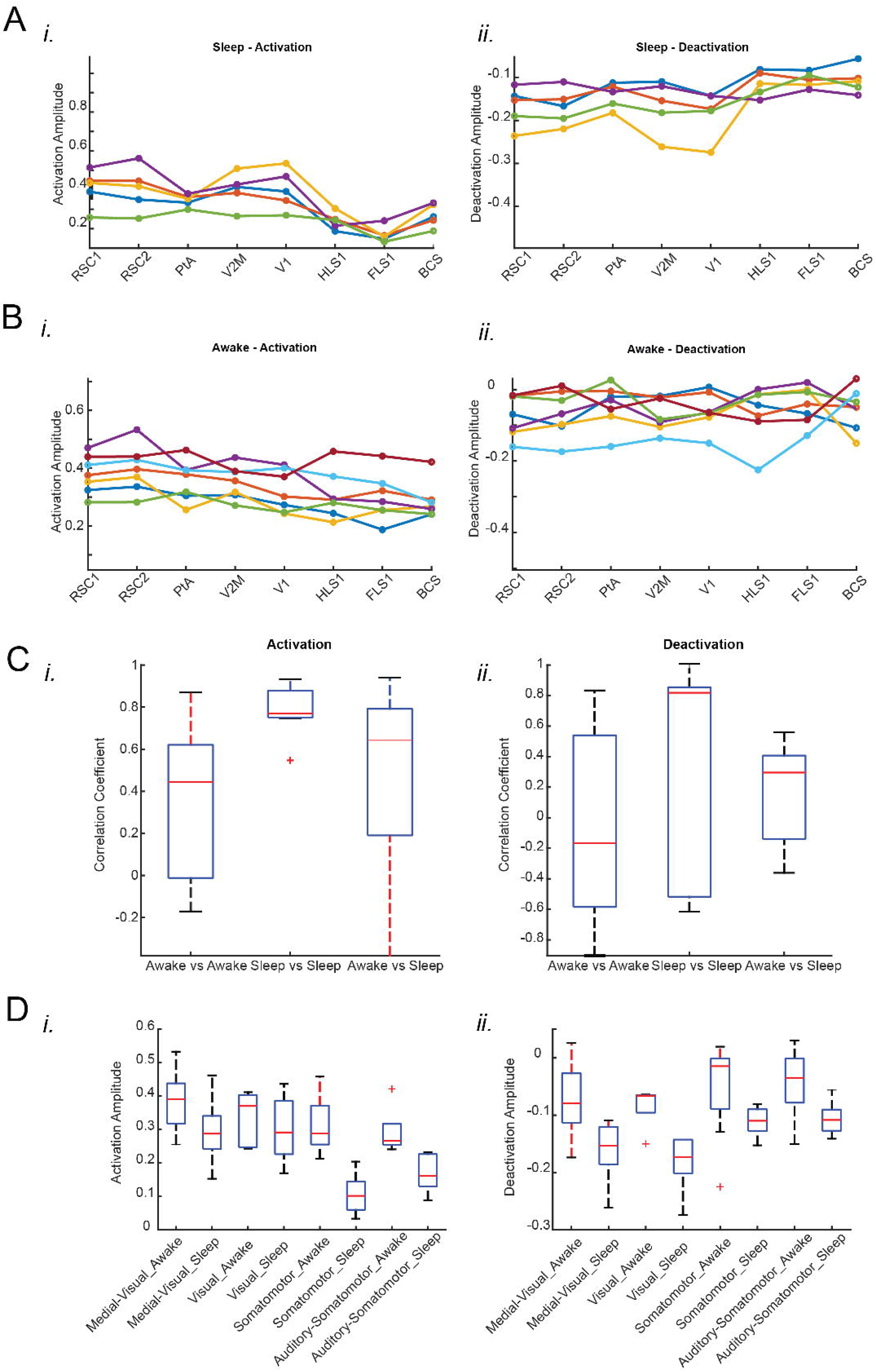

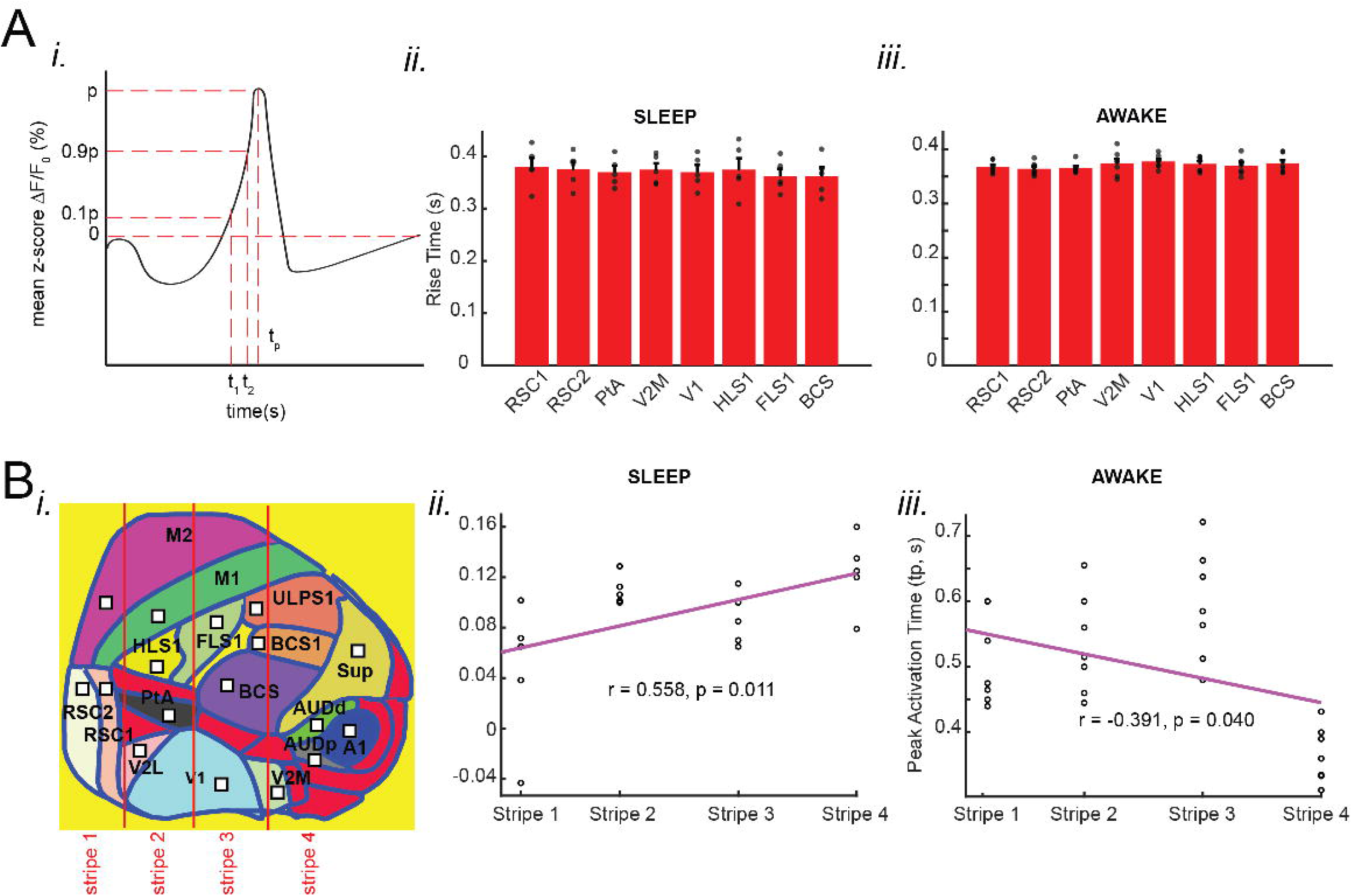

## Notes

### Competing Interest Statement

The authors have declared no competing interest.

### Summary of Updates

This version has been substantially revised throughout. The abstract, main text, analyses, figures and discussion have been updated, and the author list, affiliations and author contribution statement have been revised to reflect the contributions of additional authors.

