## Supplementary figures for "Cortical GABAergic inhibition dynamics around hippocampal sharp-wave ripples"

Supplementary Figure 1-5.

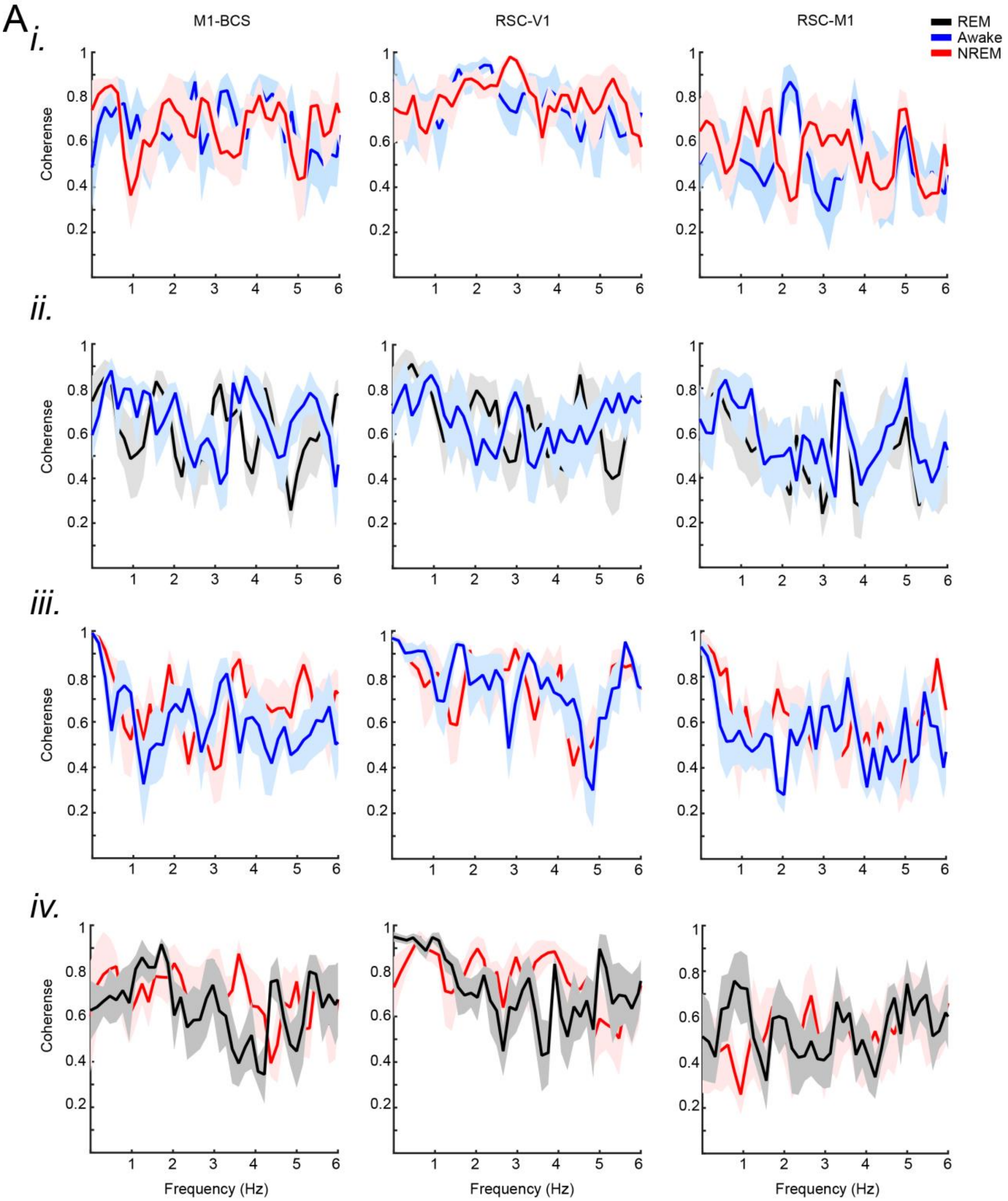


**Supplementary Figure 1. Spectral coherence analysis of iGABASnFR2 signals across dorsal cortical regions during behavioral state transitions**. Coherence spectra are shown for three region pairs (M1–BCS, RSC–V1, and RSC–M1). Panels represent transitions between states: (i) Awake → NREM, (ii) REM → Awake, (iii) NREM → Awake, and (iv) NREM → REM. Traces denote mean coherence for REM (black), Awake (blue), and NREM (red), with shaded areas indicating SEM. The analysis reveals state-dependent modulation of GABAergic inhibition and inter-regional coupling within the low-frequency range (0.5–6 Hz). Region abbreviations: M1, primary motor cortex; BCS, barrel cortex; RSC, retrosplenial cortex; V1, primary visual cortex.

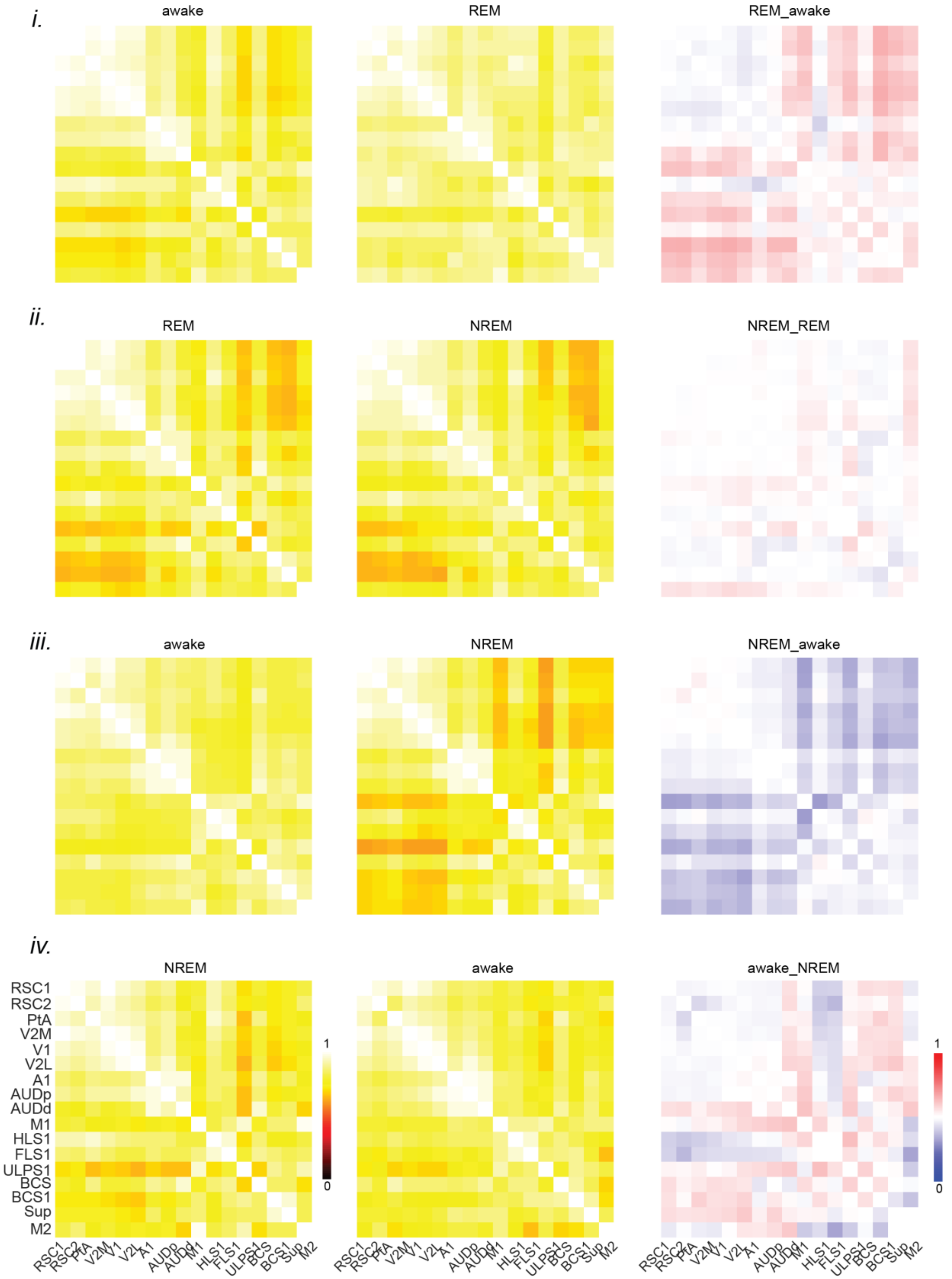


**Supplementary Figure 2. Functional connectivity matrices of iGABASnFR2 signals across dorsal cortical regions during behavioral state transitions**. Pairwise correlations are shown for

(i) Awake → REM, (ii) REM → NREM, (iii) Awake → NREM, and (iv) NREM → Awake. Each panel displays connectivity strength within a given state (left and middle) and the corresponding difference matrix between states (right). RSC1/2, retrosplenial cortex areas 1 and 2; PtA, parietal association cortex; V2M, medial secondary visual cortex; V1, primary visual cortex; V2L, lateral secondary visual cortex; A1, primary auditory cortex; AUDp/d, posterior and dorsal auditory areas; M1, primary motor cortex; HLS1/FLS1, hindlimb and forelimb primary somatosensory cortex; ULPS1, upper limb primary somatosensory cortex; BCS1/2, barrel cortex areas 1 and 2; Sup, suprasylvian cortex; M2, secondary motor cortex.

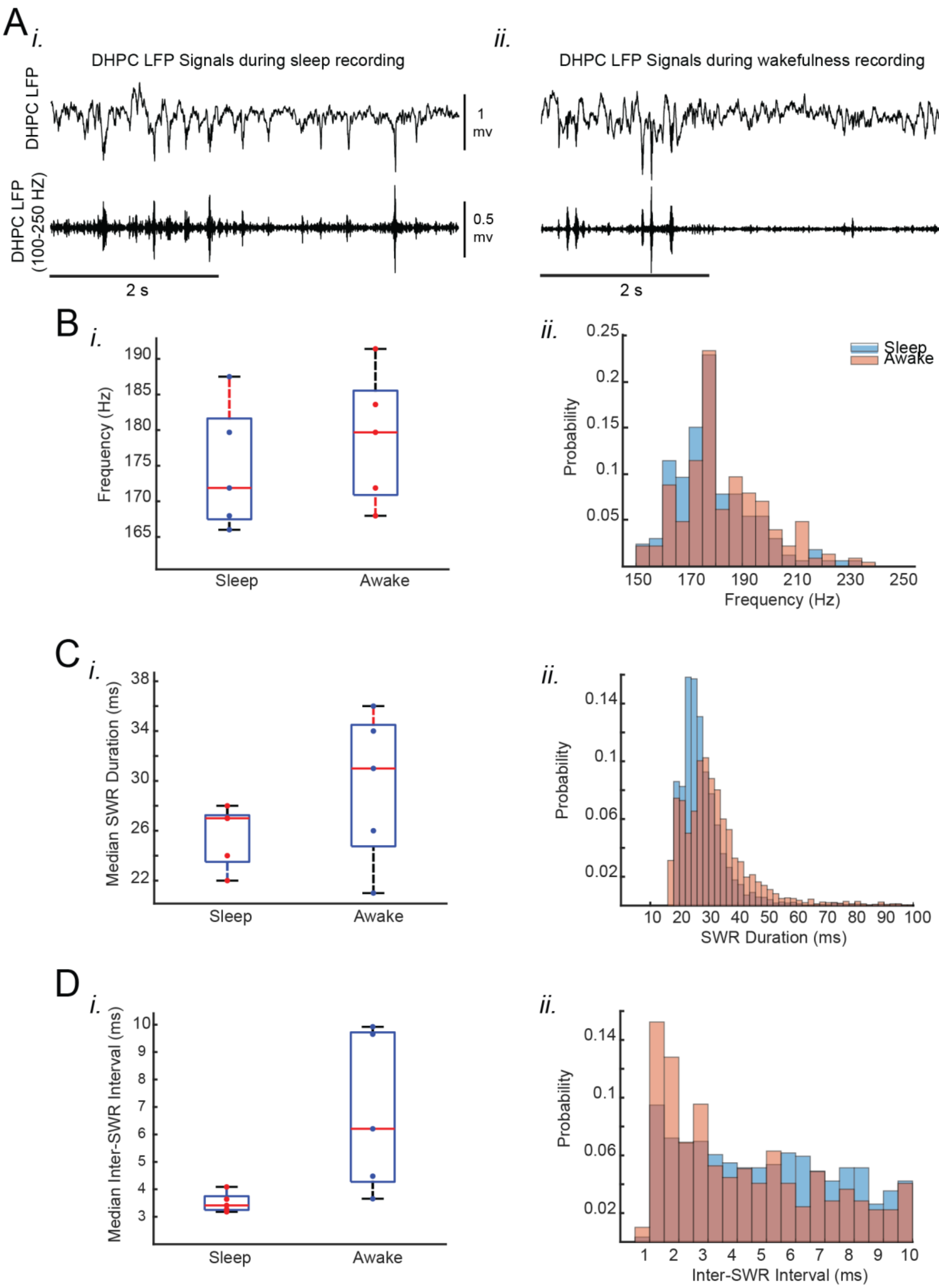


**Supplementary Figure 3. Characteristics of SWRs recorded in head-restrained naturally sleeping and wakeful mice.** (A) Representative DHPC (dorsal hippocampus) LFP signals were recorded during sleep (i) and wakefulness (ii). The top traces show raw LFP signals, and the bottom traces display band-pass filtered LFP signals (100–250 Hz). The filtered signals highlight the occurrence of sharp wave ripples (SWRs). Scale bars indicate 2 seconds. (B) (i) Median peak spectral frequency of hippocampal SWRs recorded during head-restrained sleep and wakefulness. Each data point represents the median peak frequency for a given animal (n = 7 for sleep, n = 5 for wakefulness). SWRs detected during head-restrained sleep tend to have lower peak frequencies than wakefulness. Red plus signs represent outliers beyond the interval containing 99.3% of the data points. (ii) Histograms showing the distribution of peak spectral frequencies of hippocampal SWRs pooled across all animals and experimental conditions. Sleep recordings exhibit a more clustered and lower frequency distribution, whereas wakefulness displays a broader frequency range. (C) (i) Median SWR durations during head-restrained sleep and wakefulness. Each data point represents the median value for a given animal. SWRs detected during sleep are significantly shorter than those recorded during wakefulness. (ii) Histogram distributions of SWR durations pooled across all animals and experimental conditions. Sleep recordings show a higher probability of shorter durations, whereas wakefulness is associated with more variability and longer durations. (D) (i) Median inter-SWR intervals during head-restrained sleep and wakefulness. Each data point represents the median value for a given animal. SWRs detected during sleep tend to have shorter inter-SWR intervals compared to wakefulness. (ii) Histogram distributions of inter-SWR intervals pooled across all animals and experimental conditions. Sleep recordings are associated with shorter intervals and more tightly clustered distributions, whereas wakefulness exhibits a broader range of intervals, with longer durations being more frequent.

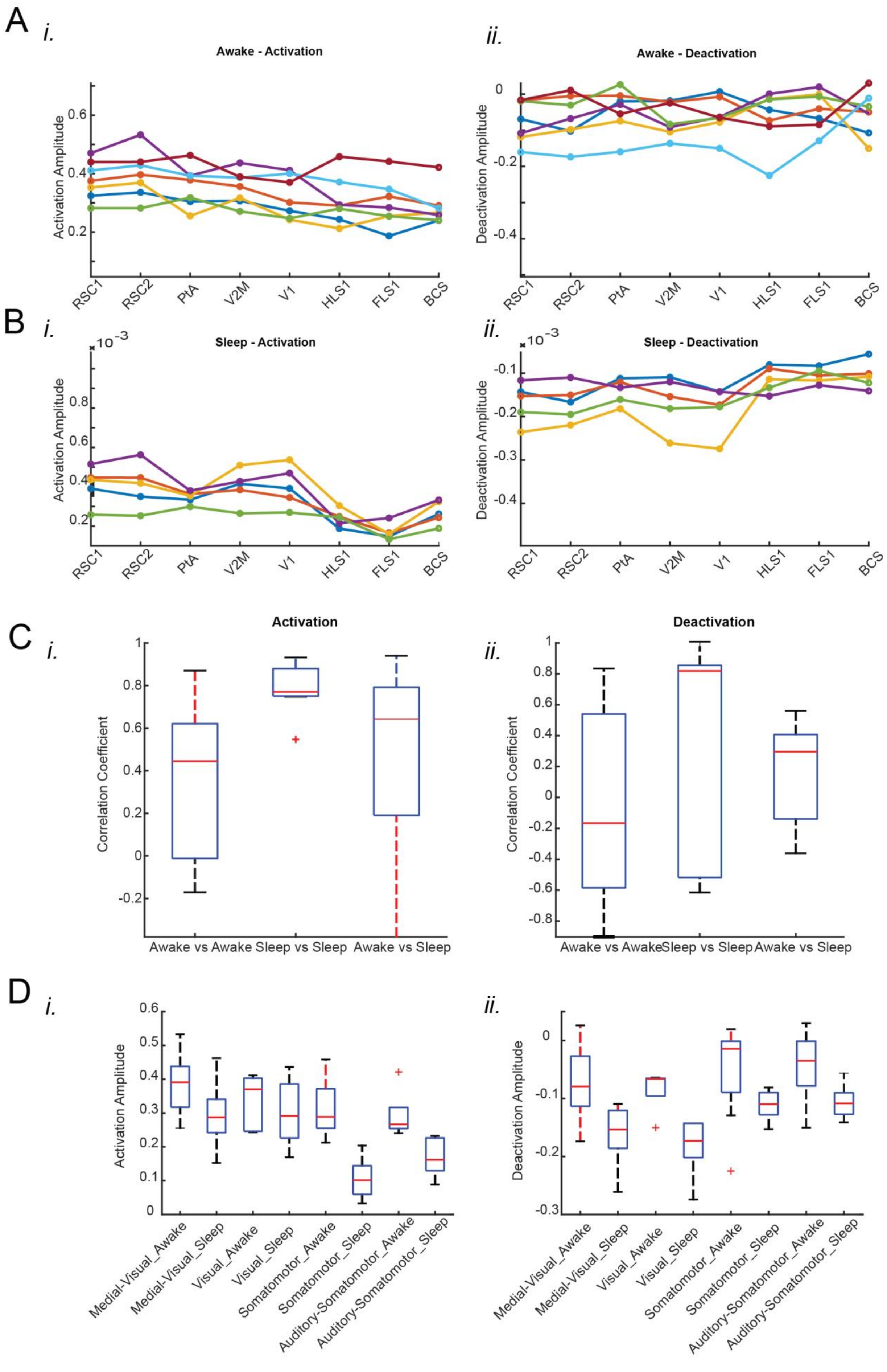

A
i.
Awake - Activation
Activation Amplitude
ii.
Awake - Deactivation
Deactivation Amplitude
RSC1
RSC2
PtA
V2M
V1
HLS1
FLS1
BCS
B
i.
Sleep - Activation
ii.
Sleep - Deactivation
x 10^-3
C
i.
Activation
Correlation Coefficient
Awake vs Awake
Sleep vs Sleep
Awake vs Sleep
ii.
Deactivation
D
i.
ii.
Medial-Visual_Awake
Medial-Visual_Sleep
Visual_Awake
Visual_Sleep
Somatomotor_Awake
Somatomotor_Sleep
Auditory-Somatomotor_Awake
Auditory-Somatomotor_Sleep

**Supplementary Figure 4. State-dependent modulation of neocortical GABA dynamics during SWRs: enhanced regional specificity in sleep compared to wakefulness.** (A) Peri-SWR GABA activation and deactivation amplitudes across distinct neocortical regions during wakefulness (i, iii) and sleep (ii, iv), sorted in decreasing order for activation in wakefulness (i) and sleep (ii). Panels (iii) and (iv) show deactivation amplitudes during wakefulness and sleep, respectively. Each piece-wise linear trace corresponds to data from an individual animal (n = 7 for wakefulness, n = 5 for sleep). The data points were averaged across regions within each neocortical structural subnetwork, and these averages were used to generate bar graphs in Figures 2Biii and 2Cii. (B) Quantification of similarities in activation (i, ii) and deactivation (iii, iv) amplitude trends across neocortical regions under wakefulness and sleep conditions. Box plots display the distribution of correlation coefficients for all possible pairs of traces for wakefulness (i, iii) and sleep (ii, iv). Horizontal red lines represent the median of each distribution, while red plus signs indicate data points outside the interval containing 99.3% of all values. Statistical analysis revealed no significant differences in trends between wakefulness and sleep, justifying the pooling of data from these two states for specific analyses. (C) Subnetwork-specific comparisons of activation (i) and deactivation (ii) amplitudes between sleep and wakefulness. The plots show the arithmetic difference in amplitudes (sleep minus wakefulness) for all possible pairs of subnetworks. A statistically significant result was not observed. These results emphasize the distinct state-dependent modulation of GABAergic dynamics, particularly in specific neocortical subnetworks.

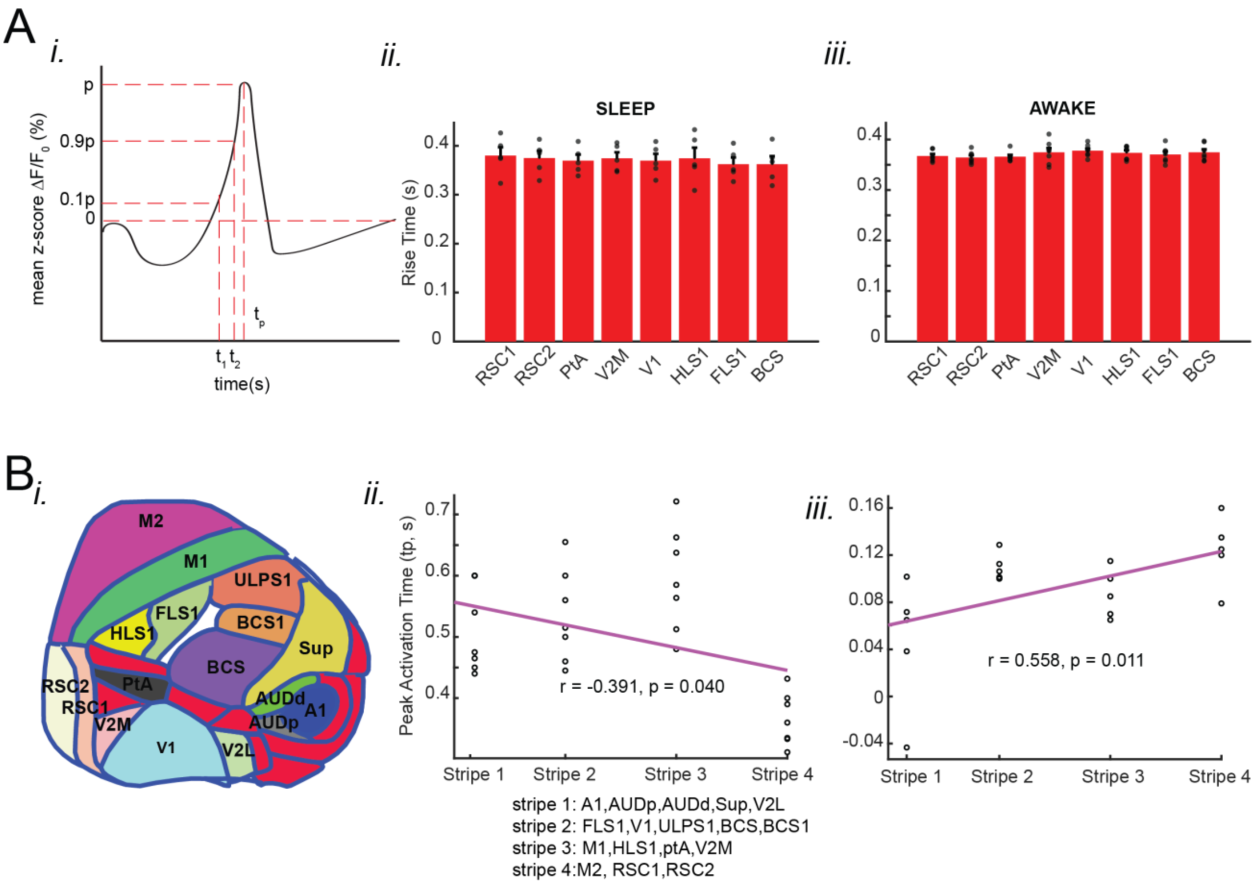

**Supplementary Figure 5. Sequential GABA Activation Across Neocortical Regions During Hippocampal SWRs: Lateral Dominance in Wakefulness and Medial Dominance in NREM Sleep (A)** GABA activation rise time in neocortical regions: (i) Measurement of GABA rise time: GABA rise time was calculated as the time taken for the signal to increase from 10% (0.1p) to 90% (0.9p) of its peak value (p). (ii) rise time during wakefulness: mean GABA rise times show no significant differences across neocortical regions. All regions activate at a similar rate. (iii) rise time during sleep: GABA rise time is also consistent across neocortical regions, with no significant regional differences. Bar graphs represent mean ± SEM, with individual data points shown. (B) sequential activation of GABA in neocortical regions relative to SWRs: (i) schematic of neocortical regions: neocortex divided into four vertical stripes from medial to lateral regions. (ii) wakefulness: A significant positive correlation ($r = 0.558$, $p = 0.011$) is observed between stripe number and peak activation time (tp), indicating that GABA activation occurs later in more lateral regions relative to hippocampal SWRs. (iii) Sleep: A significant negative correlation ($r = -0.391$, $p = 0.040$) is observed, showing that GABA activation occurs earlier in more medial regions than hippocampal SWRs.

**Video 1. Spontaneous neocortical GABA activity with simultaneously recorded hippocampal LFP and ripple activity.** This figure showcases the unique capabilities of the iGABASnFR2 sensor, capturing spontaneous neocortical GABA activity under wakefulness, filtered in the 0.5–6 Hz band, and the concurrent hippocampal local field potential recorded in two frequency bands: 0.1–1000 Hz and 100–250 Hz.

**Video 2. Wake/sleep scoring.** An example showcasing the physiological signals utilized for wake/sleep scoring during a progression from wakefulness to NREM sleep, followed by REM sleep. Notably, NREM sleep is characterized by the absence of movement, whereas REM sleep displays distinct facial muscle twitches.

**Video 3. Neocortical GABA Dynamics During NREM to Awake Transition.** This figure illustrates an example of neocortical GABA dynamics during a transition from NREM to awake, captured using iGABASnFR2 imaging. The iGABASnFR2 signal was band-pass filtered between 0.5–6 Hz to highlight the relevant frequency components. The blue traces represent the raw hippocampal LFP signal, EMG power, and the power ratio of delta to theta bands, providing additional context for the physiological changes accompanying the sleep state transition.

**Video 4. Neocortical GABA Dynamics During Awake to NREM Transition.** This figure illustrates an example of neocortical GABA dynamics during a transition from awake to NREM, captured using iGABASnFR2 imaging. The iGABASnFR2 signal was band-pass filtered between 0.5–6 Hz to highlight the relevant frequency components. The blue traces represent the raw hippocampal LFP signal, EMG power, and the power ratio of delta to theta bands, providing additional context for the physiological changes accompanying the sleep state transition.

**Video 5. Neocortical GABA Dynamics During NREM to REM Transition.** This figure illustrates an example of neocortical GABA dynamics during a transition from NREM to REM sleep, captured using iGABASnFR2 imaging. The iGABASnFR2 signal was band-pass filtered between 0.5–6 Hz to highlight the relevant frequency components. The blue traces represent the raw hippocampal LFP signal, EMG power, and the power ratio of delta to theta bands, providing additional context for the physiological changes accompanying the sleep state transition.

**Video 6. Neocortical GABA Dynamics During REM to Awake Transition.** This figure illustrates an example of neocortical GABA dynamics during a transition from REM to awake, captured using iGABASnFR2 imaging. The iGABASnFR2 signal was band-pass filtered between 0.5–6 Hz to highlight the relevant frequency components. The blue traces represent the raw hippocampal LFP signal, EMG power, and the power ratio of delta to theta bands, providing additional context for the physiological changes accompanying the sleep state transition.

**Video 7. Mean peri-SWR GABA activity during natural sleep.** An example of mean peri-SWR GABA activity captured using iGABASnFR2 imaging under natural sleep. The iGABASnFR2 signal was band-pass filtered between 0.5–6 Hz. The blue trace illustrates the mean peri-SWR hippocampal LFP signal.

**Video 8. Mean peri-SWR GABA activity during wakefulness.** An example of mean peri-SWR GABA activity captured using iGABASnFR2 imaging under wakefulness. The iGABASnFR2 signal was band-pass filtered between 0.5–6 Hz. The blue trace illustrates the mean peri-SWR hippocampal LFP signal.
